# Human subcortical pathways automatically detect collision trajectory without attention and awareness

**DOI:** 10.1101/2023.02.10.527946

**Authors:** Fanhua Guo, Jinyou Zou, Ye Wang, Boyan Fang, Huafen Zhou, Dajiang Wang, Sheng He, Peng Zhang

**Author notes:** Corresponding author (P. Zhang), (S. He) and (D. Wang). Contributed equally to this work (Fanhua Guo, Jinyou Zou, Ye Wang).

## Abstract

Detecting imminent collisions is essential for our survival and is likely supported by evolutionarily conserved mechanisms in the brain. Using high-resolution 7T fMRI, we investigated subcortical pathways for detecting collision trajectories in healthy human subjects and hemianopic patients. When healthy participants focused their attention on a central fixation task, their superior colliculus (SC), ventromedial pulvinar (vmPul) and ventral tegmental area (VTA) elicited stronger responses to a peripheral object approaching on head-collision courses compared to near-miss trajectories. Correlation and path analyses of collision-sensitive responses revealed collision sensitivity in the SC-vmPul and SC-VTA pathways without attention and cortical influence. Both behavioral performance and SC responses showed higher sensitivity to looming stimuli from the upper visual field. For hemianopic patients with unilateral lesions of the geniculostriate pathway, the ipsilesional SC, vmPul and VTA showed collision sensitivity to looming stimuli in their blind visual field, in the absence of their awareness. Stronger responses in the SC were also associated with better detection performance of the collision events. These findings clearly demonstrate that human tectofugal pathways, without attention and awareness, automatically detects approaching objects on a collision course, supporting blindsight to impending visual threats.

**Highlights:**

- SC-vmPul and SC-VTA pathways show collision sensitivity without attention and cortical influence in healthy participants.
- Both behavioral performance and SC responses show higher sensitivity to looming stimuli from the upper visual field.
- The ipsilesional SC, vmPul and VTA of hemianopic patients automatically detects collision trajectories in their blind visual field without awareness.
- SC response is associated with “blindsight” detection of impending collisions.

## Introduction

Detecting objects approaching on a collision course is critical for survival of animals in the environment. Specialized neurons or brain circuits highly sensitive to looming stimuli have been found in many species, including fruit flies, locusts, fish, frogs, pigeons and mice (Billington et al., 2011; King & Cowey, 1992; O’shea & Rowell, 1976; Salay et al., 2018; Shang et al., 2015; Y. Wang & Frost, 1992; Wei et al., 2015). Human infants also showed avoidance behaviors to symmetrically looming stimuli (Ball & Tronick, 1971). The neural mechanism for computing the time-to-contact (TTC) information of a looming stimulus has been extensively studied and was suggested to provide warning detection for large approaching objects (Fotowat & Gabbiani, 2011; Sun & Frost, 1998; Y. Wang & Frost, 1992). These previous studies mostly compared a large looming stimulus versus translating or receding stimuli, whose motion directions greatly deviate from the collision course. However, recent studies showed that humans are not only sensitive to looming vs. non-looming stimuli, but also extremely efficient in detecting collision course from near-miss trajectories. Compared with a near-miss trajectory, an approaching object on a collision course with the observer could automatically capture visual attention, increase perceived object size and evoke greater pupil constrictions, even without conscious awareness of the motion direction (Chen et al., 2016; Lin et al., 2009). These findings suggest early subcortical mechanisms for collision detection, through precise and sensitive measures of motion trajectories. However, little is known about the neural mechanism in the human brain for detection of collision trajectories.

The superior colliculus (SC) is a phylogenetically old visual nucleus lying on the roof of the mammalian brainstem. It plays important roles in visual perception and visually guided reorienting functions, such as attention, eye and head movements (May, 2006). As a key retino-recipient region, the SC and its homologous structure in non-mammals, optic tectum, were found preferentially sensitive to looming stimuli in a number of species (Billington et al., 2011; Shang et al., 2015; Wei et al., 2015; Wu et al., 2005). In rodents, specific types of neurons in the SC have been identified as the key components of several subcortical circuits to detect looming objects and trigger defensive responses, including the SC-PBGN (parabigeminal nucleus)-Amygdala (Shang et al., 2015, 2018), SC-LP (lateral posterior thalamic nucleus, homolog of primate pulvinar)-Amygdala (Shang et al., 2018; Wei et al., 2015) and SC-VTA (ventral tegmental area)-Amygdala (Zhou et al., 2019), and LC (locus coeruleus)-SC (Li et al., 2018) connections. Given that the sub-cortex is the primitive brain and that subcortical functions might be relatively conserved in mammals, the subcortical pathways found in the rodent brain might also play important roles for collision detection in the human brain. Alternatively, with the expansion of the neocortex and reduced retinal projection to the SC in higher mammals (May, 2006), it could be possible that the cerebral cortex plays a more prominent role in collision detection in the human brain. There was fMRI evidence that the human SC responded to looming but not to receding stimuli (Billington et al., 2011), in a TTC estimation task. However, in this study, only limited subcortical regions were investigated at a relatively coarse spatial scale, and it is not clear whether the SC activation was due to time-to-collision calculation or collision trajectory detection. Thus, the specific neural pathways and mechanisms for detecting collision trajectories in the human brain remain elusive.

Moreover, it is unknown whether the subcortical pathways can automatically detect imminent collisions even without attention and awareness to the looming stimuli, as suggested by previous behavioral studies (Chen et al., 2016; Lin et al., 2009). Some patients with cortical blindness can detect or even discriminate visual stimuli presented to their blind visual field, despite denial of seeing the stimuli. This phenomenon, called blindsight, attracts broad interests and has been hotly debated for almost half a century (Sanders et al., 1974; Weiskrantz et al., 1974). Although lots of studies have been done, the neural pathways involved and their specific functional roles in blindsight remain highly controversial (Cowey, 2010). Looming-evoked avoidance behavior was observed in Monkey with V1 lesions (King & Cowey, 1992), suggesting a critical role of subcortical pathways in automatic detection of impending visual threats in blindsight. However, there is no direct evidence supporting this hypothesis.

To address these questions, the current study used high-resolution fMRI to investigate the neural pathways involved in detecting collision trajectory in both healthy human participants and hemianopic patients. The motion trajectory of an incoming object was varied slightly to be either on a collision course with, or a miss trajectory of the head of observers. For healthy participants, we first measured their visual ability to discriminate collision course from near-miss trajectories and the associated changes in pupillary reflex (experiment 1). In a 7T fMRI experiment (experiment 2), we further investigated whether certain neural pathways could detect imminent collisions with or without attention. Subjects either paid attention to and judged the trajectory of the looming stimuli, or performed a central fixation task without attention to the approaching objects. Experiment 3 studied whether subcortical pathways can detect collision trajectories even without awareness and a functional geniculostriate pathway. We scanned a group of hemianopic patients with unilateral lesions of the geniculostriate pathway, while they performed a central fixation task with hit, near-miss or receding stimuli presented in their normal or blind visual fields. Four patients also measured their forced-choice detection performance to stimuli presented to their blind visual field. Our results reveal that the SC-vmPul and SC-VTA pathways in humans automatically detects collision trajectories without attention and awareness, which also supports blindsight to impending visual threats.

## Results

### Human observers can precisely discriminate collision from near-miss trajectories

In a behavioral experiment (experiment 1, n = 15), we measured the ability of healthy human observers to discriminate hit from near-miss trajectories. Subjects responded whether an incoming object would hit or miss their head (fig. 1a, fig. S1a). Pupil size and eye movements were also recorded. In fig. 1b, the percentage of hit response was plotted as a function of the extrapolated impact point of the approaching objects. The psychometric functions were further fitted with cumulative normal distributions. The steep slope of the psychometric function around the edge of the head (about 6 cm from nasion) indicates that subjects could precisely discriminate the incoming objects on a collision course from those with near-miss trajectories, consistent with the prediction from previous studies (Chen et al., 2016; Lin et al., 2009). Fig. S2a shows the results for all individuals.

**Figure 1.**
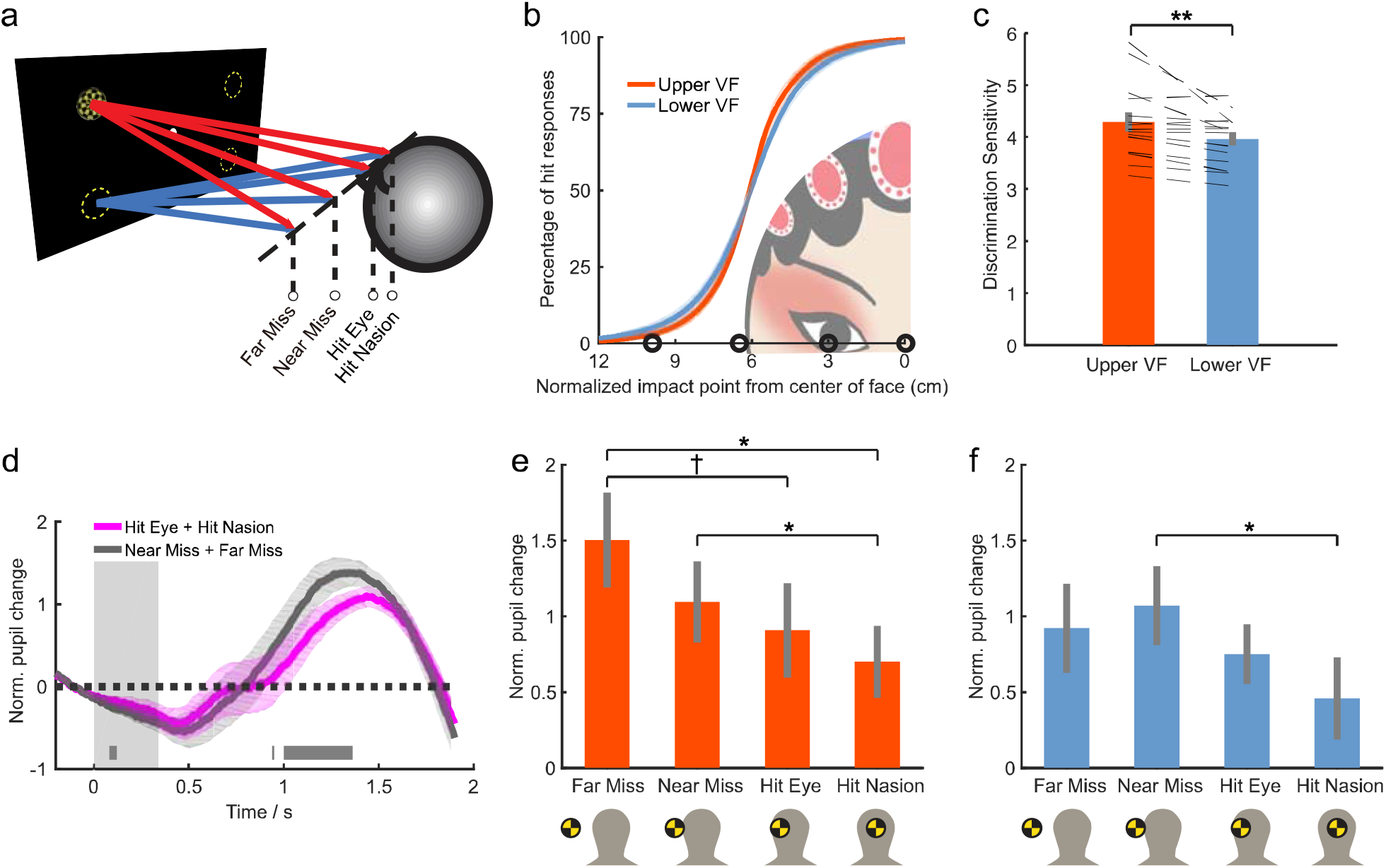
Schematic stimulus diagram and results of the behavioral experiment (exp. 1). **(a)** Visual stimuli depicting an incoming ball from one of the four quadrants of the visual field were presented with a 3D LCD monitor in the behavioral experiment. The trajectory of the looming object varied slightly to either hit (hit nasion, hit eye), or miss (near miss, far miss) the head of observers. **(b)** The percentage of hit responses at different impact points was fitted with a normal cumulative distribution function (CDF). Black circles indicate the extrapolated impact points of different trajectories. **(c)** The discrimination sensitivity, calculated as *ln*(1/*σ*) in which *σ* is the standard deviation of the fitted normal CDF, is significantly higher in the upper visual field than in the lower visual field. Error bars indicate standard error of the mean (SE). **p < 0.01. **(d)** The change of pupil size with respect to baseline (-200∼0ms) was normalized by standard deviations, and then plotted for hit (purple) and miss (dark gray) trajectories respectively (data from the upper and lower visual field were combined here). Dark gray bars indicate the time points when the pupil size was significantly smaller (permutation test p < 0.05 using cluster-size based adjustment) in the hit condition. The vertical gray bar indicates the time interval of visual stimulus presentation. **(e, f)** The averaged change of pupil size from 1000 to1364 ms after stimulus onset was plotted for different looming stimuli from the upper **(e)** and lower **(f)** visual fields. * Bonferroni corrected p < 0.05, † uncorrected p < 0.05. Shaded areas in **(b, d)** and vertical lines in **(c, e, f)** indicate SE.

The discrimination sensitivity, calculated based on the slope of the psychometric curve, was slightly but significantly higher for objects approaching from the upper visual field than from the lower visual field (permutation test p < 0.001, fig. 1c), indicating that observers could more precisely discriminate hit from near-miss trajectories for objects coming from the upper visual field. Pupil size was smaller around 1100 ms after stimulus onset in the hit than miss conditions (fig. 1d-f), consistent with previous findings (Chen et al., 2016). Fixational eye movements showed no significant difference between the hit and miss events (fig. S2b). These behavioral results show that observers and their pupillary responses can precisely discriminate collision from near-miss trajectories of an incoming object.

### Stronger responses in the SC to looming objects on a collision course compared to a near-miss trajectory

To investigate potential subcortical pathways for the detection of collision trajectory in the human brain, we performed a 7T experiment in a group of healthy human participants in exp. 2 (n = 20). In a long event-related design (fig. 2a), subjects responded whether an incoming object would hit or miss their head in an attended condition, or responded to fixation color changes irrelevant to the looming stimuli in an unattended condition.

**Figure 2.**
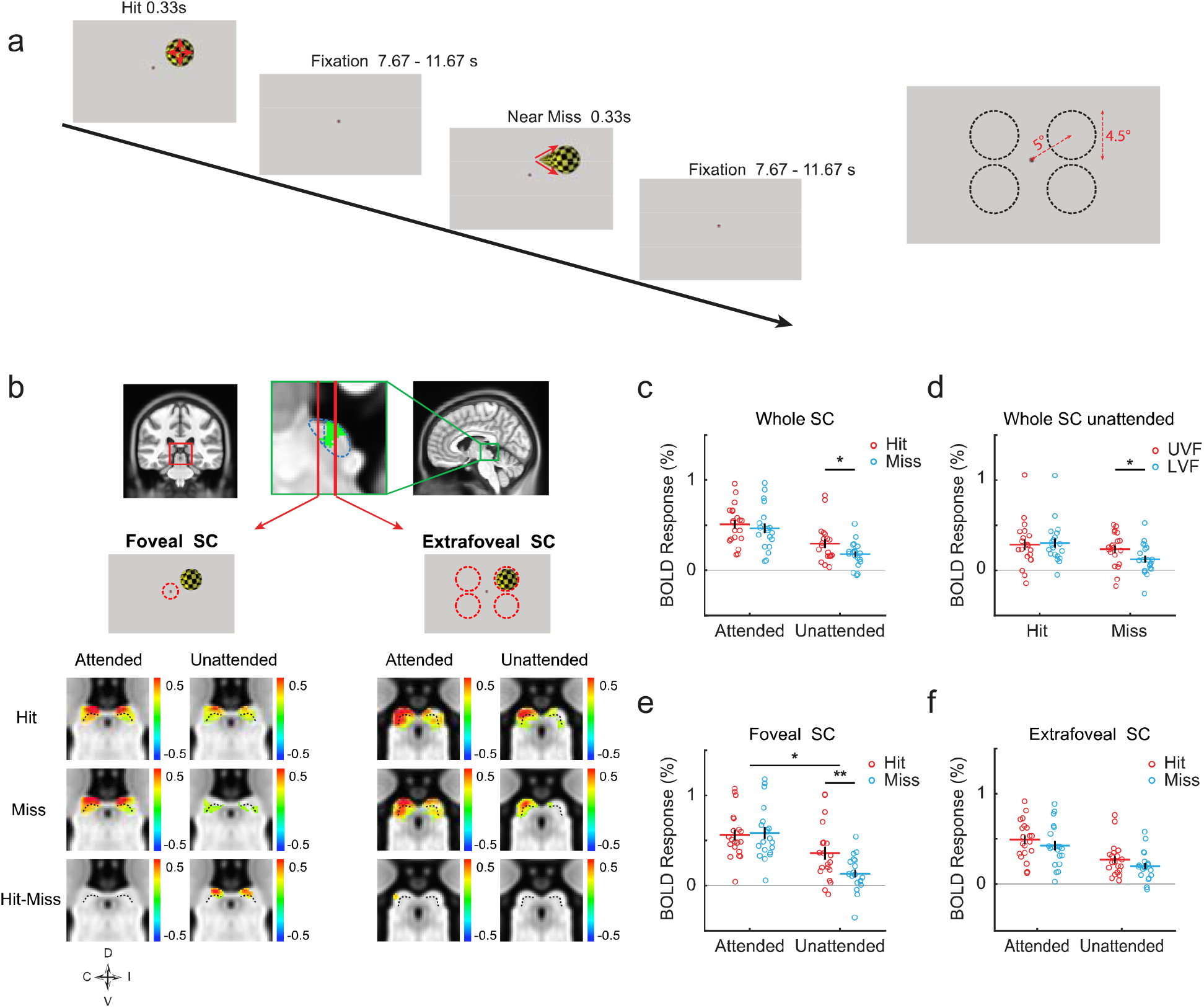
Schematic diagram of stimuli and procedure, and looming-evoked responses in the SC of healthy participants (exp. 2). **(a)** Looming objects with hit and near-miss trajectories were presented with a 2D MRI-compatible projector for 330 ms, with 7.67∼11.67 s of inter-trial intervals. The stimulus was presented in one of the four quadrants of the visual field in each trial. The right panel shows the size and position of the stimulus at the last frame in each quadrant. **(b)** SC activation maps. Red and green squares on the coronal and sagittal slices indicate the locations of the SC. From the zoomed-in sagittal view, blue dashed lines outline the ROIs of the rostral/anterior and caudal/posterior SCs representing the foveal and extrafoveal visual fields, respectively. The ROIs was determined by the retinotopic activations (contra-ipsi, fig. S4) shown here as the green overlay. Activation maps were thresholded at p < 0.05 (uncorrected). Color bars indicate percent signal change. Left and right SCs were mapped with responses to the contralateral and ipsilateral stimuli, respectively. Activation maps to stimuli from the left visual field were horizontally flipped and averaged with those to stimuli from the right visual field. C, I, D, V in the compass abbreviate contralateral, ipsilateral, dorsal and ventral, respectively. Dotted lines indicate the boundary between the superficial and deeper layers of the SC. **(c) and (d)** ROI-averaged BOLD responses of the whole SC to looming stimuli from the contralateral visual field. **(e) and (f)** show the responses in the foveal (c) and extrafoveal (f) parts of the SC. Red/blue horizontal bars, black vertical bars and red/blue circles denote mean, standard error (SE), and individual data, respectively. * above a long line indicates ANOVA p < 0.05 for the interaction between attention and trajectory. * and ** above a short line indicate paired t-test p < 0.05 and 0.01, respectively.

Previous studies suggest that the SC, a phylogenetically old midbrain nucleus, might play important roles for collision detection in the human brain (Billington et al., 2011). Thus, in the current study, we carefully investigated the SC’s responses to looming stimuli with direct-hit or near-miss trajectories. From the group-averaged activation maps (fig. 2c), bilateral SCs showed robust responses to the looming stimuli in both attended and unattended conditions. This confirmed that the human SC was indeed highly sensitive to looming stimuli. We further investigated the ***collision sensitivity*** in each voxel defined as the response difference between the direct-hit and near-miss trajectories (fig. 2c ‘Hit-Miss’). In the attended condition, no significant cluster of voxels with collision-sensitivity was found in the SC. However, significant or marginally significant clusters can be found in the unattended condition (permutation test with small volume correction: cluster’s p = 0.053 for the contralateral SC, and p < 0.001 for the ipsilateral SC. The cluster defining threshold is voxel’s p = 0.05). These collision-sensitive clusters located more rostral (or anterior) (fig. 2b, foveal SC, ‘Hit-Miss’, unattended), corresponding to the foveal part of the SC (DeSimone et al., 2015; Hafed & Chen, 2016; Savjani et al., 2018). More caudal (or posterior) part of the SC showed strong responses to both looming stimuli presented in the contralateral visual field (fig. 2c, extrafoveal SC, ‘Hit’ and ‘Miss’), but no significant collision-sensitive clusters.

The ROI analysis focused on the responses to stimuli from the ***contralateral*** visual field. We first investigated the responses of the whole SC (fig. 2c). Two-way repeated measures (rm) ANOVA revealed a significant main effect of trajectory (Hit/Miss, F(1,19) = 8.67, p = 0.008, 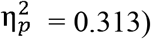) and attention (Attended/Unattended, F(1,19) = 49.27, p < 0.001, 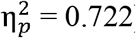). Post-hoc paired t-tests showed a significant collision sensitivity in the unattended condition (t(19) = 2.363, p = 0.029, Cohen’s d = 0.528). We then investigated the SC’s responses to looming stimuli in the upper and lower visual fields in the unattended condition (fig. 2d). Although collision sensitivity in the upper and lower visual fields showed no significant difference (trajectory by visual field interaction: p = 0.124), the SC’s responses to looming stimuli on a near-miss trajectory were significantly stronger in the upper visual field compared to those in the lower visual field (t(19) = 2.798, p = 0.011, Cohen’s d = 0.626), suggesting higher looming sensitivity in the SC to approaching objects from the upper visual field. This finding is consistent with the behavioral results of slightly better looming-trajectory discrimination and stronger pupillary reflex in the upper visual field (fig. 1).

We further divided the SC into a rostral (foveal) part and a caudal (extrafoveal) part (enclosed by blue dotted lines in fig. 2b) based on the significant contralateral-ipsilateral activations (green voxels in the upper middle panel of fig. 2b, see also fig. S4). In the foveal SC (fig. 2e), there was a significant interaction between trajectory and attention (F(1,19) = 8.06, p = 0.010, 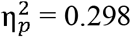). Post-hoc t-tests showed significant collision sensitivity in the foveal SC in the unattended condition (t(19) = 3.015, p = 0.007, Cohen’s d = 0.674), but not in the attended condition (p = 0.600). To further validate the collision sensitivity found in the foveal SC in the unattended condition, we performed a leave-one-subject-out (LOSO) cross-validation analysis. For each subject, the ROI to calculate the collision-sensitive response was defined by voxels with significant group-level collision sensitivity from the remaining subjects. The LOSO results revealed a significant collision sensitivity in the foveal SC in the unattended condition (t(19) = 2.761, p = 0.012, Cohen’s d = 0.617). In the extrafoveal SC (fig. 2f), there was a significant main effect of trajectory (F(1,19) = 6.75, p = 0.018, 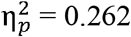).

### Collision sensitive responses in the ventral pulvinar and VTA

We further investigated collision sensitivity in other subcortical regions, including the pulvinar, VTA, PBGN, amygdala and LC, which form looming sensitive circuits with the SC as shown by previous rodent studies (Li et al., 2018; Shang et al., 2015, 2018; Wei et al., 2015; Zhou et al., 2019). The lateral geniculate nucleus (LGN) was also included as a potential control area. Fig. S3 shows the anatomical masks for these subcortical regions in MNI space. For each ROI, we performed similar analyses as for the SC to check its activation map and ROI-averaged responses. Collision sensitive responses were found in the pulvinar and VTA (fig. 3), but not in other subcortical nuclei (fig. S5).

**Figure 3.**
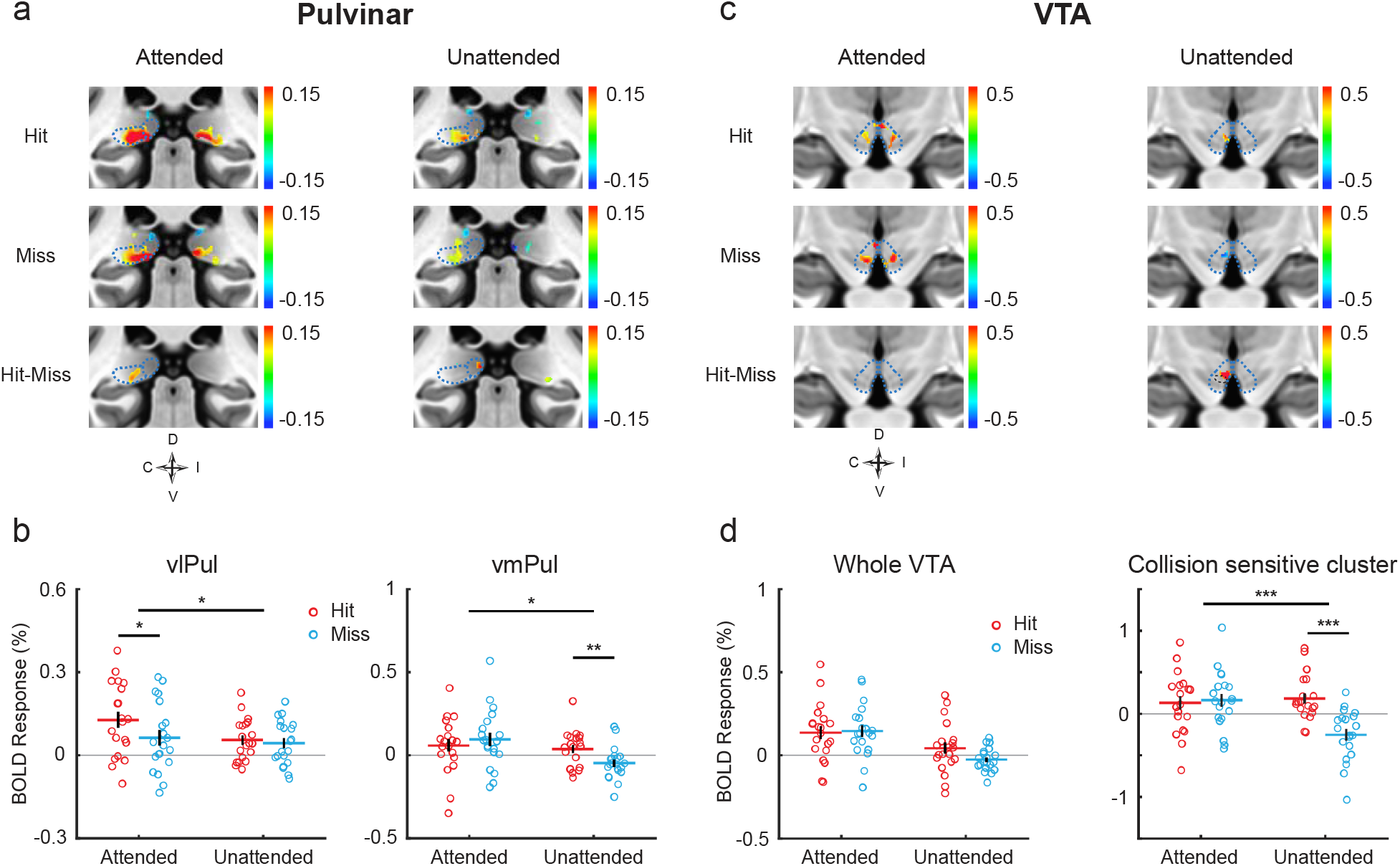
BOLD responses to looming stimuli in the pulvinar and VTA (exp. 2). **(a, c)** Activation maps to looming stimuli in coronal views of the pulvinar **(a)** and VTA **(c)**. Maps were thresholded at p < 0.05 uncorrected. Dotted lines mark the anatomical boundaries of the vlPul, vmPul and VTA. **(b)** ROI-averaged responses of the vlPul and vmPul. **(d)** Left panel shows the ROI-averaged responses of the whole VTA. Right panel shows the averaged responses of the collision sensitive cluster in **(c)**. * or *** above a long line indicate ANOVA p < 0.05 or p < 0.001. *, **or *** above a short line for t-test p < 0.05 or p < 0.001. Other conventions as in fig. 2.

As shown by fig. 3a, looming stimuli activated both lateral and medial portions of the ventral pulvinar. The ventrolateral pulvinar (vlPul) mainly connects with the early visual cortex but also with the parietal cortex (Arcaro et al., 2015, 2018), and may also receive sparse input from the SC (Lyon et al., 2010; Stepniewska, 1999). As shown by fig. 3a, a marginally significant collision sensitive cluster was found in the vlPul in the attended condition (cluster’s p = 0.091). In fig. 3b (left panel), the ROI-averaged responses of vlPul to stimuli presented to the contralateral visual field showed significant interaction of attention and trajectory (F(1,19) = 4.62, p = 0.045, 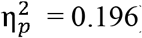). Post-hoc paired t-tests revealed a significant collision sensitivity in the attended condition (t(19) = 2.750, p = 0.013, Cohen’s d = 0.615), but not in the unattended condition (p = 0.347). The LOSO cross-validation analysis revealed a significant collision sensitivity in the vlPul in the attended condition (t(19) = 2.646, p = 0.016, Cohen’s d = 0.592).

The ventromedial pulvinar (vmPul) receives strong inputs from the SC and possibly sparse input from the retina, reciprocally connects with visual areas in the dorsal visual stream (Berman & Wurtz, 2010; Bridge et al., 2016; Kaas & Lyon, 2007; Stepniewska, 1999). The dorsal part of vmPul may also connect with the amygdala and frontoparietal cortex (Jones & Burton, 1976; Romanski et al., 1997). From the group-averaged activation maps in the unattended condition (fig. 3a), there was a small cluster of collision sensitivity (fig. 3a bottom right). The ROI-averaged responses of vmPul (fig. 3b right panel) showed a significant interaction between trajectory and attention (F(1,19) = 7.06, p = 0.016, 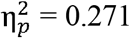). Post-hoc t-tests revealed a significant collision sensitivity in the unattended condition (t(19) = 3.613, p = 0.002, Cohen’s d = 0.808), but not in the attended condition (p = 0.321). LOSO cross-subjects validation didn’t find significant effect of collision sensitivity, likely due to a relatively large inter-individual variability of collision sensitive response in the vmPul. No significant collision sensitive response was found in other sub-nuclei of pulvinar.

The VTA receives direct input from the SC (Coizet et al., 2003; Comoli et al., 2003; Dommett et al., 2005; Zhou et al., 2019), and projects to many brain areas including the amygdala (Tang et al., 2020; Zhou et al., 2019) and frontal lobe (Beier et al., 2015; Coenen et al., 2018). In the attended condition (fig. 3c left), the VTA showed significant activations to looming stimuli but without collision-sensitivity. In the unattended condition, there was a significant collision sensitive cluster (cluster’s p = 0.004). The averaged responses of the collision sensitive cluster were plotted in the right panel of fig. 3d. For the ROI-averaged response of the whole VTA (fig. 3d, left panel), we found a marginally significant effect of collision sensitivity in the unattended condition (t(19) = 1.970, p = 0.064, Cohen’s d = 0.441). LOSO analysis further revealed a significant collision sensitivity in the unattended condition (t(19) = 3.234, Cohen’s d = 0.723, p = 0.004).

### Collision-sensitive responses in the visual cortex and frontoparietal cortex

We further investigated whether collision sensitivity can also be found in the cortical brain areas. The retinotopic responses of most responsive voxels in the visual cortex (see methods for ROI definitions) were shown in fig. 4a. Significant collision sensitivity can be found in the early visual cortex in both attended and unattended conditions (Attended condition: q < 0.001 in V1 with BH-FDR correction, q = 0.011 in V2, q = 0.006 in V3, q = 0.012 in V4; Unattended condition: q = 0.003 in V1, q = 0.006 in V3, q = 0.007 in V4; uncorrected p = 0.027 in TO2). Fig. 4b shows the group-averaged collision-sensitive (Hit-Miss) activations on the cortical surface. A few clusters with collision sensitivity (p < 0.01, uncorrected) can be found from bilateral middle temporal gyrus (MTG) in the attended condition, and from bilateral temporal parietal junctions (TPJ), and the frontal eye field (FEF), inferior frontal gyrus (IFG) and insular (INS) in the right hemisphere in the unattended condition. However, no significant cluster can be found after family-wise error correction. These results revealed collision-sensitive responses in the visual cortex and possibly in the frontoparietal areas of attention network.

**Figure 4.**
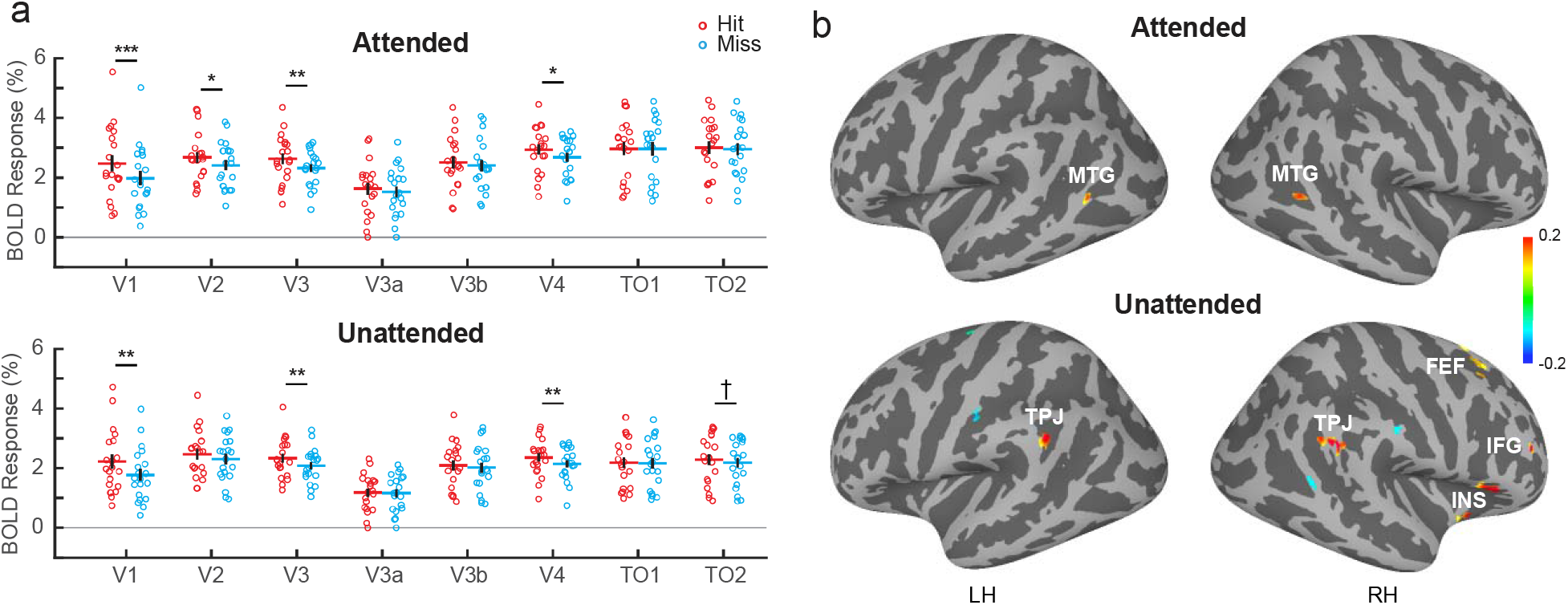
Collision sensitivity in the cortical areas (exp. 2). **(a)** BOLD responses in the visual cortex. *, ** or *** denote q < 0.05, q < 0.01 or q < 0.001 after FDR correction, and † for p < 0.05 (uncorrected). **(b)** Collision-sensitive activations in the high order visual cortex and the frontoparietal areas. The color bar indicates the percent signal change of Hit-Miss. Maps were thresholded at p < 0.01 (uncorrected).

### SC-vmPul and SC-VTA pathways detect collision trajectories without top-down attention and cortical influence

To further investigate the subcortical circuits involved in collision detection and the potential influence from cortical brain areas, we calculated across-subject correlations between collision-sensitive responses in the subcortical nuclei (SC, vmPul and VTA) and cortical areas including the visual cortex (VC) and frontoparietal attention network (AttNet), followed by a path-analysis with structural equation modeling (SEM) to infer their causal relationships. We didn’t include vlPul in this analysis because the SC’s projection to vlPul is weak and controversial (Lyon et al., 2010; Stepniewska, 1999), and the collision sensitive responses in the SC and vlPul showed no significant correlation (p>0.49 in both attended and unattended conditions).

In the attended condition (fig. 5a, left panel), we found significant correlations of collision sensitive responses between cortical and subcortical regions (SC-VC: Spearman’s r (rs) = 0.606, p = 0.006; SC-AttNet: rs = 0.606, p = 0.006; vmPul-VC: rs = 0.731, p < 0.001), and also between the SC and its downstream subcortical nuclei (SC-vmPul: rs = 0.474, p = 0.036; SC-VTA: rs = 0.484, p = 0.032). In the unattended condition, there were significant correlations between the SC and its downstream targets (SC-vmPul: rs = 0.570, p = 0.010; SC-VTA: rs = 0.654, p = 0.002), but not between cortical and subcortical regions (fig. 5a, right panel, all p > 0.19). Here we used Spearman’s correlation because there are outliers in the data pairs of the correlation matrix.

**Figure 5.**
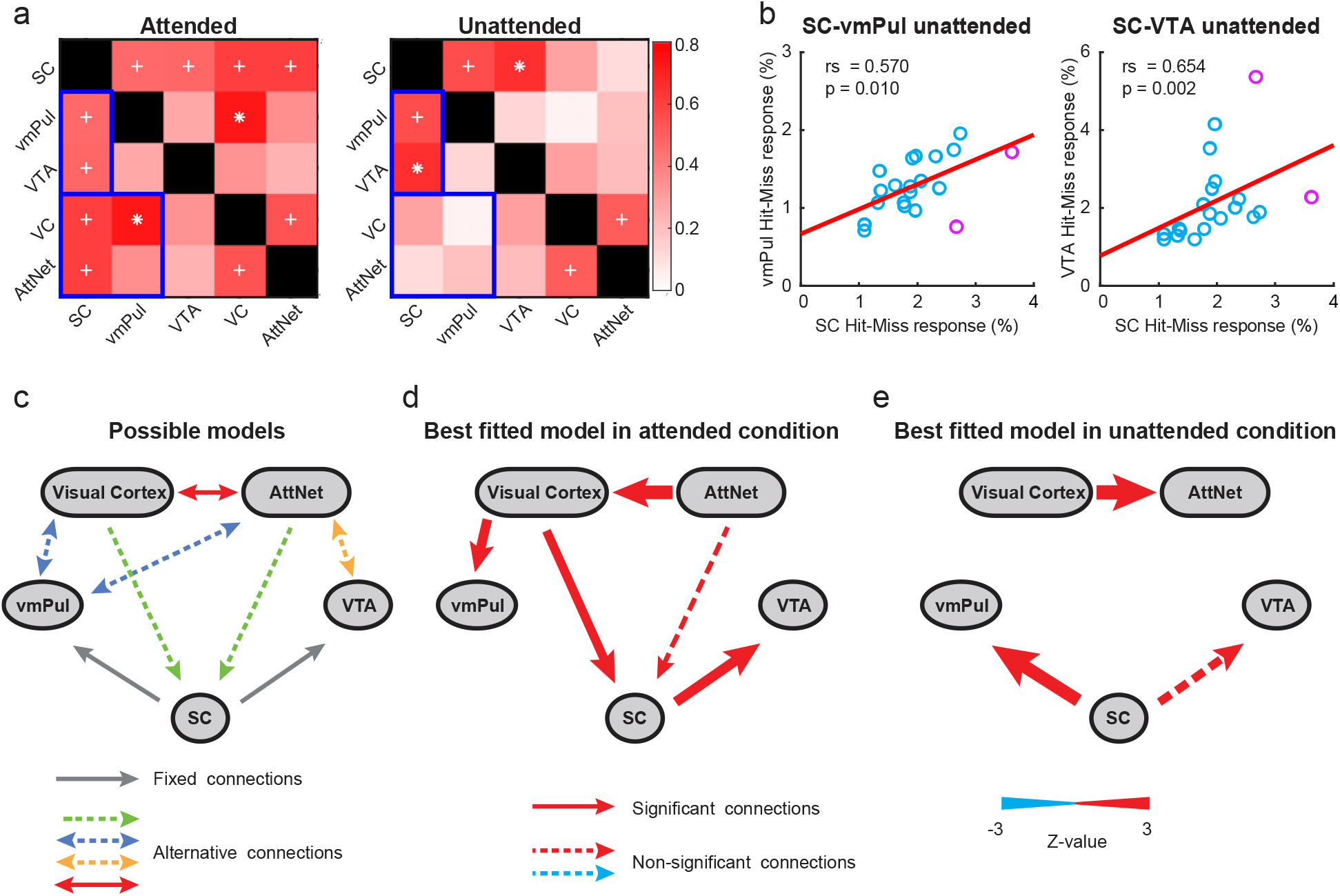
Correlation and path analyses of collision sensitive responses between cortical and subcortical areas (exp. 2). **(a)** Correlation matrix. Each grid in the matrix shows the Spearman’s correlation coefficient between the collision sensitivities of two ROIs, in the attended (left) and unattended (right) conditions. + indicates uncorrected p < 0.05 and * for p<0.05 after FWE correction by permutation test. Colorbars indicate the correlation coefficient. Blue squares indicate significant correlations between SC and vmPul in both attended and unattended conditions, whereas significant correlations between subcortical and cortical areas only in the attended condition. **(b)** Scatterplots of individual data showing the correlations between collision sensitivity SC-vmPul and SC-VTA. **(c)** Candidate SEM models for the causal relationships between ROIs. Gray solid single-headed arrows represent fixed one-way connection. Red solid double-headed arrows indicate connections with two alternative directions. Dotted double-headed arrows indicate that the connection may follow one of two alternative directions or may not exist. All combinations of alternative connections yield 216 candidate models. **(d, e)** Best fitted models in the attended **(d)** and unattended **(e)** conditions. Arrows with solid lines indicate significant connections. Arrows with dashed lines indicate non-significant connections. Arrow thickness denotes the z-value of connections. Insignificant connections with p > 0.5 were not shown.

To identify the collision-detection pathways from the SC and the causal influence from cortical areas, we performed a SEM path-analysis based on the correlations of collision-sensitive responses between these ROIs. 216 candidate models were constructed with different combinations of 6 sets of alternative connections (fig. 5c), based on the known anatomical connections between these areas: SC-cortex (May, 2006), SC-vmPul (Kaas & Lyon, 2007), SC-VTA (Zhou et al., 2019), vmPul-cortex (Arcaro et al., 2018), VTA-cortex (Beier et al., 2015; Zubair et al., 2021). Candidate models were fitted with the observed data and compared by the goodness of fit (see Methods for model comparisons). The best-fitted model for the attended condition (fig. 5d, fit index: *χ*2 = 0.075, df = 2, CFI = 1.000, GFI = 0.998, AGFI = 0.989, RMSEA = 0.000, RMR = 0.010) revealed significant connections from the visual cortex to the SC (p = 0.043) and vmPul (p = 0.031), and from SC to VTA (p = 0.025). In the unattended condition, the best fitted model (fig. 5e, fit index: *χ*2 = 0.157, df = 2, CFI = 1.000, GFI = 0.997, AGFI = 0.977, RMSEA = 0.000, RMR = 0.020) revealed a robust SC-vmPul connection (p = 0.004) and a marginally significant SC-VTA connection (p = 0.056), but without significant cortical influence (all p > 0.5). Using a beta-series method (Cisler et al., 2014; Rissman et al., 2004), there were indeed significant trial-based functional connectivity of SC-vmPul (t(19) = 3.260, Cohen’s d = 0.729, p = 0.004) and SC-VTA (t(19) = 4.651, Cohen’s d = 1.040, p < 0.001) in the unattended condition. These results support a pivotal role of the SC-vmPul and SC-VTA pathways in detecting collision trajectories without attention and cortical influence.

### The ipsilesional tectofugal pathways of hemianopic patients are sensitive to collision stimuli in their blind visual field

To further investigate whether the tectofugal pathways can detect collision trajectories even without awareness and a functional geniculostriate pathway, we scanned a group of homonymous hemianopic patients with unilateral lesions of the geniculostriate pathway (experiment 3, n = 12). Patients lost their conscious vision of both eyes in one side of the visual field (see table S1 for clinical characteristics, and fig. S7 for visual field loss and lesioned locations). During fMRI scans, patients performed a central fixation task while stimuli with hit, near-miss, or receding trajectories were presented to their normal visual field (NVF) or blind visual field (BVF). Four patients (P09∼P12) also participated in a behavioral visibility test. In which, they reported clear perception of stimuli presented to their NVF but denied seeing stimuli from the BVF. For most patients (8 of 12), there was no significant V1 activation to stimuli presented in their BVF (fig. S7). Four patients showed weak uncorrected activations in their lesioned hemisphere, but no collision sensitivity.

Based on the findings of normal participants in exp. 2, we focused our analysis on the SC, vmPul and VTA in hemianopia patients. The LGN was also investigated since previous studies suggest that it plays a critical role in blindsight (Schmid et al., 2010). From the group-averaged activation maps in fig. 6a, significant clusters of activation to hit trajectories were found in the contralateral SCs regardless of whether the stimuli were presented in the NVF (cluster’s p = 0.002) or in the BVF (cluster’s p = 0.030). Also, there was no significant difference between hit responses in the ipsilesional and contralesional clusters (100 most responsive voxels were selected from each cluster: permutation test p = 0.155. We used permutation test in patient data due to the relatively small number of subjects and the presence of outliers.). In contrast, there was no significant activation to near-miss or receding stimuli in both contralateral SCs. In the ipsilesional SC, a small cluster of voxels with collision sensitivity was found to looming stimuli presented in the BVF.

**Figure 6.**
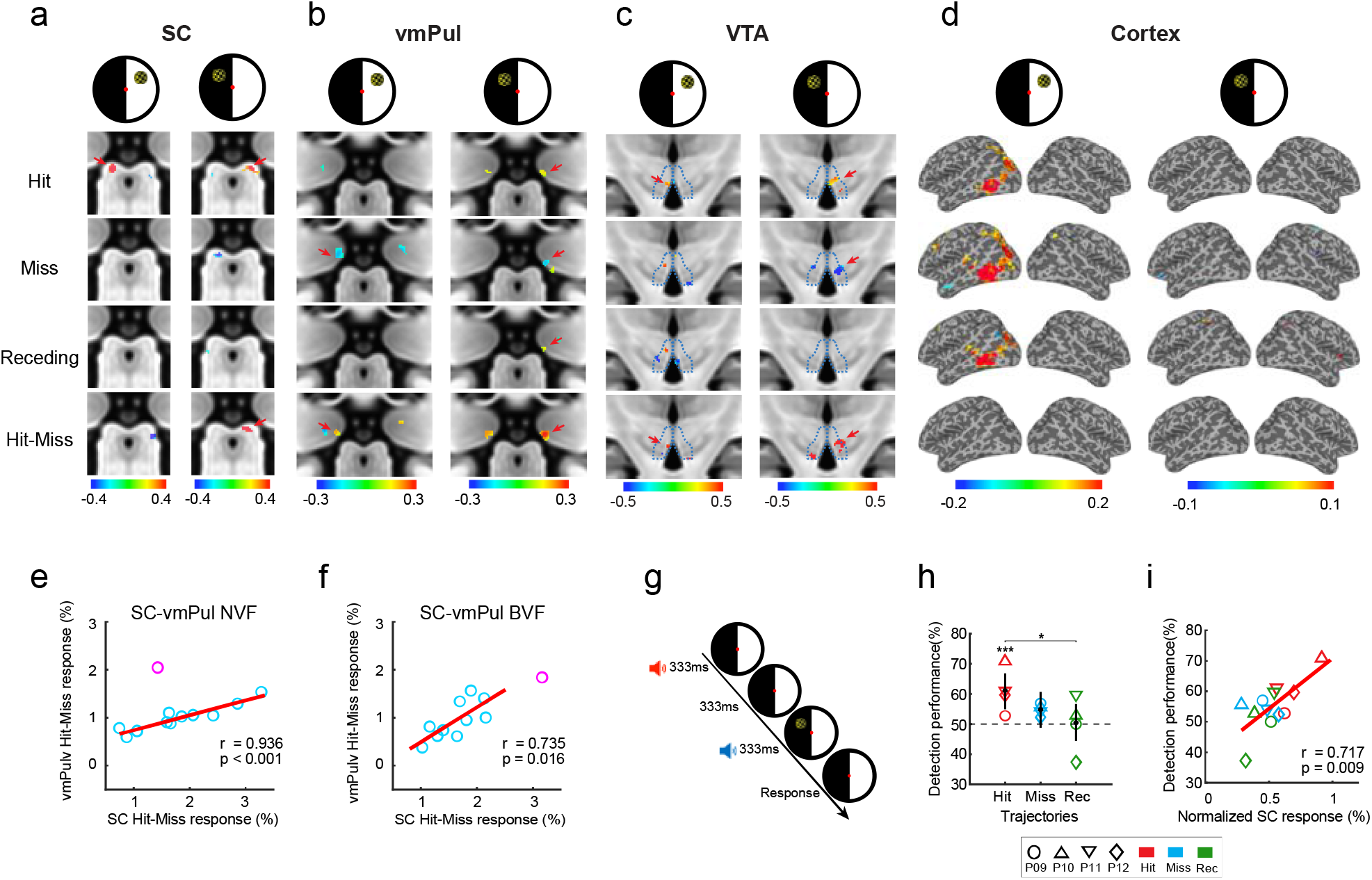
FMRI results and behavioral performance of hemianopic patients (exp. 3). **(a)** Group-averaged activation maps in the SC to stimuli presented to the NVF (left column) and BVF (right column). Maps were thresholded at p < 0.05 uncorrected. Color bars represent percent BOLD signal change. The left and right SCs were mapped contralateral to the NVF and BVF, respectively. **(b, c)** Activation maps in the vmPul (b) and VTA (c). Conventions are the same as in (a). Only activations within the ROI were shown. Red arrows indicate the location of collision-sensitive clusters from the Hit-Miss map. **(d)** Stimulus activations mapped to the cortical surface (thresholded at p < 0.01 uncorrected). Left columns show contra- and ipsi-lateral responses to stimuli presented to the NVF. Right columns show ipsi- and contra-lateral responses to stimuli presented to the BVF. **(e, f)** Correlations between collision sensitive responses in the SC and vmPul contralateral to the NVF (e) and BVF (f). Each circle represents one patient. Red circles indicate the outliers. **(g)** Schematic diagram and procedure for the 2-IFC detection task. **(h)** The detection permanence fitted with a binomial generalized linear mixed model. Error bars represent the 95% confidence intervals. * post-hoc test p < 0.05 for comparing marginal means between conditions. *** p < 0.001 for comparing marginal means to the chance level (50%). **(i)** The correlation between the detection performance to stimuli in the BVF with the ipsilesional SC’s responses (in 10% most responsive voxels, chosen separately for each stimulus type). For each subject, SC responses were normalized by dividing the mean across three stimulus conditions and then multiplying the mean across all four subjects.

For the vmPul, the group-averaged activation maps in fig. 6b revealed a significant collision sensitive cluster in the ipsilesional pulvinar to stimulus presented in the BVF (cluster’s p = 0.026), and a smaller cluster at a similar location in the contralesional pulvinar when stimuli were presented in the NVF. LOSO analysis demonstrated significant collision sensitivity in the ipsilesional pulvinar to looming stimuli in the BVF (permutation test p = 0.008). Moreover, individually defined collision sensitivity (strongest Hit-Miss responses in each ROI, see methods for details) showed a significant correlation between the contralateral SC and vmPul when stimuli were presented in the BVF (rs = 0.782, p = 0.007, fig. 6g), and a marginally significant correlation when stimuli were presented in the NVF (rs = 0.539, p = 0.075, fig. 6f).

A significant collision sensitive cluster was also found in the ipsilesional VTA to stimuli in the BVF (cluster’s p = 0.042). Collision sensitive responses in the VTA showed an insignificant trend of correlation with those in the SC (NVF: rs = 0.483, p = 0.115, BVF: rs = 0.413, p = 0.185). No collision-sensitive response was found in the LGN (fig. S8). In the cortical regions, V1 (fig. S7) and frontoparietal areas (fig. 6d) showed strong activations to stimuli presented in the NVF but not in the BVF. These findings clearly demonstrate that the tectofugal pathways can automatically detect collision trajectories without awareness and a functional geniculostriate pathway.

### Stronger response in the ipsilesional SC is associated with better detection performance of collision events in the blind visual field

To check potential “blindsight” to approaching objects on a collision course, four patients (P09∼P12) also performed a two-interval forced-choice (2-IFC) detection task. The stimulus was presented in one of two 330-ms intervals with a 330-ms gap in-between, accompanied by a low-pitch and a high-pitch tones in the first and the second intervals, respectively (fig. 6g). Patients had to determine in which interval the stimulus was presented. The detection performance was fitted with a binomial generalized linear mixed model (fig. 6h). Results showed a significant main effect of trajectory (*χ*^2^ (2) = 6.418, p = 0.04). Following tests showed that the accuracy for the object on a collision course was significantly higher than that on a receding trajectory (contrast = 10.6%, z = 2.529, p = 0.023 using Holm adjustment), and than chance level, i.e., 50% (estimated marginal mean = 61.1%, z = 3.541, p < 0.001). Importantly, the detection performance significantly correlated with the response of ipsilesional SC (Pearson’s r = 0.717, p = 0.009; Main effect of SC responses on detection performance in a linear mixed model: F(1,3.05) = 14.616, p = 0.031; fig. 6d, right panel). No significant correlation was found between behavioral performance and the response of vmPul, probably due to the lower signals from this region (fig. 6i). Altogether, the behavioral and fMRI results of hemianopic patients in exp. 3 provide strong evidence in humans that the tectofugal pathways support blindsight to detect impending collisions.

## Discussion

Detecting imminent collision is crucial for our survival. In exp. 1, we found that human observers can precisely discriminate whether an approaching object was on a collision course or a near-miss trajectory with their head. Collision events also induced significant changes in pupillary reflex. In exp. 2, high-resolution 7T fMRI revealed collision-sensitive responses in several subcortical nuclei, including the SC, ventral pulvinar and VTA. Correlation and path analyses further demonstrated collision sensitivities in the SC-vmPul and SC-VTA pathways without attention and cortical influence. In exp. 3, for hemianopic patients with unilateral lesions of the geniculostriate pathway, the ipsilesional SC, vmPul and VTA showed collision sensitivity to stimuli presented to their blind visual field. Finally, stronger response in the SC was associated with better detection performance of the collision event. These findings clearly demonstrate a critical role of the human tectofugal pathways in automatic detection of collision trajectories without attention and awareness, supporting “blindsight” to threating visual information.

### Human SC is highly sensitive to the collision trajectory of a looming object

In the optic tectum and the downstream nucleus rotundus (homologues of the SC and pulvinar in mammals) of pigeons, different types of neurons encode several optical variables of a looming stimulus, including the time-to-collision, absolute rate of expansion and object size (Sun & Frost, 1998; Y. Wang & Frost, 1992; Wu et al., 2005). In the mouse SC, Shang and colleagues identified parvalbumin-positive (PV+) excitatory projection neurons in the superficial layers encoding the optical parameters of a looming stimulus in their receptive fields (Shang et al., 2015, 2018). The response onset depended on the stimulus size and moving velocity, and the response peaked at the time of collision. Here we show that the human SC was highly sensitive to slight trajectory differences of looming stimuli with similar times to contact, rates of expansion and object sizes, suggesting that the primate SC contains neurons sharply tuned to looming trajectories for accurate collision detection. Interestingly, although the looming stimuli were presented in the parafovea, the strongest collision-sensitive response was observed in the anterior part of the SCs corresponding to the central visual field (fig. 2c). This finding may suggest that SC neurons are sensitive to the would-be point of collision, making predictions about the impact point in the near future.

The observed collision sensitive response in the SC was unlikely due to fixational eye movements, because observers can maintain stable fixations that showed no significant difference in fixational eye movements between the hit and miss conditions (fig. S1b). The lack of collision sensitivity in the attended condition was likely due to a saturation effect of the SC response.

### Upper visual field advantage of sensitivity to looming stimuli

In exp.1, the behavioral performance and pupillary reflex can better discriminate looming trajectories from the upper visual field (fig. 1). The SC response was also stronger to looming stimuli from the upper visual field (fig. 2d). This upper visual field advantage of looming sensitivity suggests that the phylogenetically conserved tectal pathways are also ecologically adaptive, as threatening looming objects such as diving predators or falling stones appear more often in the upper than in the lower visual field due to gravity. A recent study showed that the primate SC, like that of rodents, also over-represents the upper visual field, with sharper, stronger and faster visual representations than in the lower visual field (Hafed & Chen, 2016). Together, these findings support Previc’s ecological perspective of the primate visual system, suggesting that the primate SC may be optimized and specialized for detection of transient and biological salient information from extra-personal space in the upper visual field (Previc, 1990). This upper visual field advantage forms an interesting contrast with previous findings that sustained visual attention resolves and tracks objects better in the lower visual field (He et al., 1996).

### Attention but not awareness strongly modulates looming sensitive response in the SC

The results of exp.2 clearly show that compared with the looming-evoked responses during a central fixation task, paying attention to and judging the trajectory of looming stimuli strongly enhanced the responses in both superficial and deeper layers of the SC (fig. 2, fig. S6c). However, in exp.3, the SC’s responses to collision trajectories were comparable when the stimulus was presented to the normal visual field or to the blind visual field of hemianopic patients (fig. 6a). Thus looming sensitive responses in the SC were strongly modulated by attention to but not awareness of the looming stimuli. These findings demonstrate the critical role of the SC in both top-down attention and subconscious processing of threatening visual information, supporting the important claim that top-down attention and consciousness are two distinct processes of the brain (Cohen et al., 2012; Koch & Tsuchiya, 2007).

### Human tectofugal pathways automatically detect collision trajectories without attention and awareness

Our data clearly demonstrate that the SC-vmPul pathway in the human brain automatically detects collision trajectories without attention and awareness to the looming stimuli. This finding is consistent with recent rodent studies that the SC-LP (or pulvinar) pathway processes looming information in an anesthetized state (Shang et al., 2018) and triggers defensive freezing behavior (Shang et al., 2018; Wei et al., 2015). In Wei et al’s study, optogenetic mapping and electrophysiology also revealed a di-synaptic circuit from the SC through LP to the lateral amygdala, which directly mediated the innate fear-related defensive response. In our study, although looming objects on a collision course changed pupillary reflex, subjects didn’t report fear to these stimuli, and no significant response was found in the amygdala (fig. S5). One possible explanation is that human observers quickly adapted to repetitive presentations of the virtual looming objects, which may not be effective to trigger fear response or evoke strong activation in the amygdala. In contrast, collision sensitive responses can be reliably observed in the SC and vmPul. We speculate that the role of the SC-vmPul pathway might be automatically processing collision-related visual information, such as the looming trajectory or time-to-collision, which could be used by the downstream brain areas (e.g. the dorsal stream) to guide quick and subconscious actions to avoid the impending threats (Goodale, 1993).

Collision-sensitive response was also found in the VTA without attention and awareness (fig. 3c, d and fig. 6b), showing correlation with and effective connectivity from the SC (fig. 5b, e). These findings are consistent with the rodent studies that VTA neurons responded in short latency to biologically salient or aversive stimuli (Coizet et al., 2003; Comoli et al., 2003; Guarraci & Kapp, 1999), and that VTA neurons receiving direct input from the SC mediated defensive flight behaviors to large overhead looming stimuli (Zhou et al., 2019). Populated mainly with dopaminergic (DA) neurons, the VTA plays important roles in reward, motivation and attention (Schultz, 1998; Schultz & Dickinson, 2000). The role of SC-VTA pathway might be modulating the level of arousal (Eban-Rothschild et al., 2016) or changing the attentional state (Redgrave et al., 1999; Richter & Gruber, 2018) of the observer to mediate rapid defensive responses (Becerra et al., 2001).

Although rodent studies also suggest the PBGN and LC in processing looming information and mediating looming-evoked defensive behaviors (Li et al., 2018; Shang et al., 2015), we didn’t find significant collision-sensitive response from these areas in the human brain. Given their small sizes and deep locations, the negative finding from these small subcortical nuclei could be due to the low SNR and partial volume effect of BOLD signals.

### The tectofugal pathways support ‘blindsight’ detection of impending visual threat

The tectopulvinar pathway was originally proposed to support “blindsight” (Weiskrantz et al., 1974). A critical role of the SC in visuomotor functions has been repeatedly confirmed by studies of visually guided saccades in V1-lesioned monkeys (Kato et al., 2011; Mohler & Wurtz, 1977; Rodman et al., 1990; Solomon et al., 1981). FMRI studies of blindsight patients also indicated visually evoked activation from the SC (Morris et al., 2001; Sahraie et al., 1997; Tamietto et al., 2010; Tamietto & de Gelder, 2010). However, given the small sample sizes and the low-resolution fMRI approaches in these studies, the involvement of the SC in human blindsight still lacks conclusive evidence. Using high-resolution fMRI, here we showed clear evidence from 12 hemianopic patients that visually evoked response in the ipsilesional SC was highly sensitive to looming objects on a collision course (fig. 6a), which also predicted the above-chance detection performance of impending collisions (fig. 6d). These findings provide strong evidence for the critical role of the SC in detecting impending visual threats in human blindsight.

Compared with the SC, the role of pulvinar in blindsight is more controversial. Human fMRI studies suggest that the tecto-pulvinar-amygdala pathway may underlie blindsight of emotionally and socially salient face stimuli (Ajina et al., 2020; de Gelder et al., 2005; Morris et al., 2001). A recent study in V1-lesioned monkeys also revealed a critical role of the tectopulvinar pathway in visually guided saccades (Kinoshita et al., 2019; Takakuwa et al., 2021). However, there is accumulating evidence that the geniculo-extrastriate pathway plays a major role in blindsight in both Monkey (Schmid et al., 2010; Takakuwa et al., 2021; Yu et al., 2018) and Human (Ajina et al., 2015; Ajina & Bridge, 2018). These discrepancies might depend on the scope of lesion, time of lesion, or distinct roles of the pulvinar and LGN pathways in blindsight (Isa & Yoshida, 2021; Rima & Christoph Schmid, 2020; Tamietto & Morrone, 2016). Our data clearly support the last hypothesis. Collision sensitive response was found in the ipsilesional ventromedial pulvinar to looming stimuli presented to the BVF (fig. 6b,e), which showed strong correlations with the SC (fig. 6f). While visually evoked responses can be found in the LGNs of both healthy participants (fig. S6) and hemianopic patients (fig. S8), there was no collision sensitivity. In addition to the SC-vmPul pathway, collision sensitive response was also found in the VTA (fig. 6c), suggesting that the SC-VTA pathway was also involved in ‘blindsight’ detection of threatening information. These findings support different roles of the subcortical pathways in blindsight: the tectofugal pathways are specialized to detect impending visual threats, while the geniculo-extrastriate pathway processes basic visual information (e.g. contrast and motion).

### Implications for a simple and rapid neural computation of collision trajectory

A few human fMRI studies reported cortical representations of the TTC and possibly trajectories of approaching objects (Field and Wann 2005, Coull, Vidal et al. 2008, Calabro, Beardsley et al. 2019). While the cortical responses may indicate a complex and relatively slow neural calculation for 3D motion perception (Vanduffel, Fize D Fau - Peuskens et al., Duhamel, Colby et al. 1998, Katsuyama, Usui et al. 2011), the subcortical pathway found in our study suggest a simple and rapid computational strategy for collision trajectory detection. In support of this, studies have demonstrated that simpler equations for calculating looming trajectories could better predict human performance, especially the systematic errors made by observers (Harris and Drga 2005, Duke and Rushton 2012). For example, (Duke and Rushton 2012) suggested that the perceived trajectory is based on the ratio of lateral angular speed to the sum of looming and changing disparity signals. Our findings revealed the neural substrate for a simple and rapid computation of collision trajectory in the human brain, which may guide the optimization of computer vision algorithms for collision detection under time pressure, e.g., (Shigang and Rind 2006).

## Acknowledgement

This study was funded by National Science and Technology Innovation 2030 Major Program (2022ZD0211900, 2021ZD0204200), National Natural Science Foundation of China (31871107, 31930053), Strategy Priority Research Program (XDB32020200) and Key Research Program of Frontier Sciences of the Chinese Academy of Science (KJZD-SW-L08), Beijing Natural Science Foundation (7212092) and the Capital’s Funds for Health Improvement and Research (2022-2-5041).

## Methods and Materials

### Experiment 1: behavior and eye tracking in healthy participants

#### Participants

15 healthy subjects (20 to 30 years old, mean age = 24.6 years, SD = 2.16 years, 10 females) participated in experiment 1. They had normal or corrected-to-normal vision without neurological or psychiatric disorders. All participants (including those for experiment 2 and 3) gave written informed consent in accordance with procedures and protocols approved by the Institutional Review Board of the Institute of Biophysics, Chinese Academy of Sciences.

#### Stimuli and procedures

Visual stimuli were generated with Psychtoolbox 3.0 in MATLAB (Brainard, 1997; Pelli, 1997). 3D rendered spheres depicted from slightly different perspectives were projected into two eyes to produce a three-dimensional effect. Stimuli were stereoscopically presented with shutter glasses (NVIDIA 3D VISION) and a compatible LCD display with 120Hz refresh rate (60Hz for each eye). Eye positions and pupil size of the left eye were recorded at 1000 Hz with an Eyelink1000Plus system. A standard 9-point calibration was performed at the beginning of each session. A low-pass filter with cut-off frequency of 40 Hz was performed to the eye-tracking data to reduce the 60 Hz artifacts from shutter glasses. After interpolating the missing data due to eye blinks, pupil-size and eye-position time series were epoched and averaged across trials per condition and subject.

As shown in fig. S1a, the 3D display presented a baseball sized sphere in 6-cm diameter moving at a speed of 24 m/s from 11.3 meters to 3.3 meters in front of the observer. The trajectory started from one of the four quadrants at a 0.65-meter horizontal offset and a 0.38-meter vertical offset, and ended in the same quadrant. The would-be point of collision varied from the middle of the face (0 cm, hit nasion) to 3 cm (hit eye), 6 cm (near miss) or 12 cm (far miss) of horizontal offset. To match their retinotopic locations, the on-screen projections of looming trajectories were aligned based on their center of mass. The on-screen display was a sphere with a texture of black-and-yellow checkerboard expanding from 0.3 to 1 degree of visual angle in 330 ms, at an eccentricity of 3.8 degrees (angle with vertical meridian is 60º). Subjects were instructed to keep fixation and press buttons to report whether the looming object would hit or miss their head. 400 trials were collected for each subject.

### Experiment 2: fMRI study with healthy participants

#### Participants

20 healthy subjects (22 to 42 years old, mean age = 26.3 years, SD = 4.0 years, 11 females) with normal or corrected-to-normal vision and no neurological or psychiatric conditions participated in experiment 2.

#### Stimuli and procedures

Visual stimuli were rendered in 3D but presented with an MRI safe 2D projector on a translucent screen behind the head coil. Participants viewed the stimuli through a mirror mounted inside the head coil. Looming objects (fig. S1b) were simulated to approach from 8.75 m from the observer and vanish at 0.75 m, at a speed of 24 m/s. The ball would either hit the eye of the observer to cause a collision (hit) or slightly miss the head (near miss) with a 6-cm horizontal offset. Same as in experiment 1, the on-screen projections of the two trajectories in the same quadrant were aligned based on their center of mass. The stimulus on the screen expanded from 0.4 to 4.5 degrees of visual angle in 330 ms, at an eccentricity of about 5 degrees (angle with vertical meridian is 60º).

In the attended condition, subjects were instructed to keep fixation and respond whether the incoming object was on or off a collision course with their heads. In the unattended condition, they were instructed to keep fixation and detect color changes of the fixation point. Subjects performed 4 runs each for the attended and unattended conditions. Each run comprised 32 trials with an inter-stimulus-interval (ISI) randomly chosen from 8, 10 and 12 seconds. Thus, 16 trials were collected for each combination of attention, trajectory and quadrant conditions. Subject S19 lost one trial of data in some conditions due to a technical problem.

#### MRI data acquisition

MRI data were acquired with a 7T scanner (Siemens MAGNETOM, Erlangen, Germany) with a 32-channel receive 1-channel transmit head coil (Nova Medical, Cambridge, MA, USA), at Beijing MRI center for Brain Research (BMCBR). The gradient coil has a maximum amplitude of 70 mT/m, 200 us minimum gradient rise time, and 200 T/m/s maximum slew rate. Functional images were acquired with a T2*-weighted 2D GE-EPI sequence (1.5-mm in-plane resolution, 1.5-mm slice thickness without gap, 68 axial slices, TR = 2000 ms, TE = 21.6 ms, flip angle = 80°, image matrix = 122×122, FOV = 183×183 mm, partial Fourier factor = 6/8, bandwidth = 1576 Hz/Px, GRAPPA acceleration factor = 2, phase encoding direction from A to P). A few EPI images with reversed phase encoding direction (P to A) were also acquired to correct image distortions in the phase encoding direction. Anatomical images were acquired with a T1-weighted MP2RAGE sequence (0.7-mm isotropic voxels, 256 sagittal slices, FOV = 224×224 mm, TR = 4000 ms, TE = 3.05 ms, TI1 = 750 ms, flip angle = 4º, TI2 = 2500 ms, flip angle = 5º, bandwidth = 240 Hz/Px, phase partial Fourier = 7/8, slice partial Fourier = 7/8, GRAPPA = 3). To improve data quality, a bite-bar was used to restrict head motion.

#### MRI data preprocessing

MRI data were analyzed using AFNI (Cox, 1996), FreeSurfer (version 6.0) (Dale et al., 1999; Reuter et al., 2012), ANTs (Avants et al., 2011), and the lab-developed mripy package (https://github.com/herrlich10/mripy). The preprocessing of volumetric data includes slice timing correction, EPI image nonlinear distortion correction with reversed-blip method, linear motion correction, T1w anatomical image registration to EPI volumes, spatial normalization to MNI space, and per-run scaling to percent signal change before GLM analysis. For spatial normalization of subcortical areas, we estimated a 12-parameter linear transformation from the anatomical volume to a high-resolution MNI template (ICBM 152 2009c symmetric T1w template), followed by nonlinear transformation with ANTs (v2.1.0) using a subcortical mask focused on the areas of interest. All spatial transformations (including motion correction, anatomical to functional volume registration, and spatial normalizations) were combined altogether and applied to the functional volumes in one interpolation step (cubic method) at 0.6 mm isotropic resolution (J. Wang et al., 2022).

A surface-based approach was used for cortical data analysis. The T1w MP2RAGE anatomical volume was segmented into white matter, gray matter, and cerebrospinal fluid using FreeSurfer’s automated procedure, using its high-resolution option. The preprocessed volumetric data before spatial normalization were mapped to the inflated cortical surface. Surface data were then normalized to a standard surface (std141) with surface-based warping, followed by surface-based smoothing with a 4.5mm FWHM (full-width half-maximum) gaussian kernel (Argall et al., 2006). Surface data were then normalized to percent signal change before GLM analysis.

#### General linear model (GLM) analysis

BOLD signal change from baseline for each stimulus condition were estimated using a GLM with fixed HRFs. For cortical data, we used a canonical HRF (BLOCK4 in AFNI). For subcortical data, we used a faster and narrower HRF that is more appropriate for subcortical regions (Lewis et al., 2018). Motion parameters, their derivatives and square of derivatives, were included as regressors of no interest.

#### ROI definition and group-level statistical maps

As in fig. S3, ROIs for the SC, PBGN, LC, VTA, and LGN were carefully drawn on the MNI template according to their anatomical landmarks and atlases (Ding et al., 2016; Mai et al., 2015; Paxinos et al., 2012). ROIs for the pulvinar subdivisions were obtained from a parcellation based on task-coactivation profiles (Barron et al., 2015). The ROI of amygdala was defined based on a manually delineated high-resolution atlas of subcortical nuclei (Pauli et al., 2018). Before group-level analysis, spatial smoothing was performed on individual beta maps of GLM results *within* the ROI with a 2.83 mm FWHM Gaussian filter for the pulvinar and amygdala, and 1.4 mm FWHM for other subcortical nuclei. Group-level statistical maps were then generated using standard random-effects analysis (t-test) on the beta maps across subjects.

Cortical ROIs were defined on the cortical surface. ROIs of visual cortical areas were defined with a retinotopic atlas generated with the Human Connectome Project retinotopy dataset (Benson et al., 2014, 2018). Frontoparietal ROIs were defined on the standard surface by the group-level activation of the mean response across all stimulus conditions (fig. S7, p < 0.05, FWE corrected).

#### ROI-averaged responses

For the subcortical ROI, we obtained the averaged response of the whole ROI as defined by fig. S3. To validate the collision sensitivity in a subcortical area with independent voxel selection, a leave-one-subject-out (LOSO) cross-validation approach was used. For each subject, the ROI to calculate the collision-sensitive response was defined by voxels with group-level collision sensitivity (hit-miss, p < 0.05 uncorrected) from the remaining subjects. For the visual cortex, the retinotopic ROI corresponding to the stimulus location in each quadrant was defined as significantly activated voxels across all stimulus conditions for each individual (p < 0.05 uncorrected). The top 10% of most significantly activated voxels were selected based on the T value of mean response. The selected voxels from all quadrants were then pooled together to calculate the ROI-averaged responses for the visual cortical area.

#### Correlation analysis of collision sensitive responses

To investigate the relationship between different brain areas in collision detection, we calculated Spearman’s correlations between the collision-sensitive responses in subcortical and cortical areas across subjects. We used Spearman’s correlation because of the outliers in data pairs. In this analysis, voxels with the strongest collision sensitive responses were selected for each ROI and for each individual. Given the relatively high SNR in the SC, we selected the top 10% of voxels with the strongest activations. For all other ROIs, the top 20% of voxels were selected to calculate collision sensitivity. Nevertheless, using top 20% voxels in the SC gave similar results. For cortical areas, we selected the top 1% voxels of collision sensitivity.

#### Causal inference by structural equation modeling

Path analysis under SEM framework was performed to infer the causal influence of collision-sensitive responses between subcortical regions (SC, vlPul, vmPul and VTA), visual cortex (including V1, V2, V3, V3a, V3b and TO1/TO2) and frontoparietal attention networks (including IPS, FEF, TPJ and IFJ). The collision-sensitive responses across all subjects were concatenated as the model input. 216 possible models were constructed by all combinations of 6 groups of possible alternative connections. By doing so, we could directly compare the strength and possibility of our alternative hypotheses. In addition, several common connections were added in all models based on the prior knowledge. All path coefficients in each model were freely determined by maximum likelihood estimate in the SEM with lavaan software 0.6-3 (Rosseel, 2012). The model selection was based on several indices. We first excluded models with an adjusted goodness-of-fit index (AGFI) less than 0.1, then ranked the remaining models by their parsimony goodness-of-fit index (PGFI) and finally picked the best fitted models for the attended and unattended conditions (JC Zhuang et al., 2005). Other goodness-of-fit indices including *χ*2, comparative fit indices (CFI), goodness-of-fit index (GFI), root mean square residual (RMR) and root mean square error of approximation (RMSEA) were also calculated and ranked to ensure the best model was not sensitive to the ranking methods. Different ranking methods led to the same results.

#### Beta-series functional connectivity

A beta-series method was adopted to test whether functional connectivity existed between the SC and vmPul (Cisler et al., 2014; Rissman et al., 2004). The response for each trial was estimated with a separate regressor in a GLM analysis of the ROI-averaged BOLD time series, generating a series of beta values for each ROI. The beta series for each trajectory condition were Z-scored to avoid the influence of mean amplitude differences. A second-level linear regression was conducted with any two series of these beta-values from two different ROIs. And the second-level regression coefficient was obtained as the functional connectivity.

### Experiment 3: fMRI study with hemianopic patients

#### Participants

12 hemianopic patients (23 to 64 years old, mean age = 44.8 years, SD = 13.8 years, 1 female) with unilateral lesions of the geniculostriate pathway, and no other neurological or psychiatric conditions were enrolled from General Hospital of People’s Liberation Army, Beijing, China. Patients lost their conscious vision in one half or a quadrant of the visual field from both eyes. They had normal or corrected-to-normal vision outside the blind visual field. Clinical characteristics of all patients were shown in table S1. Their Humphrey perimetry and lesioned locations were shown in fig. S8. The scotoma was defined as the visual field with a relative sensitivity less than -20 dB and p < 0.5% compared with normal population in both eyes.

#### Stimuli and procedures

Visual stimuli and procedures were similar as those in experiment 2. Looming trajectories were slightly changed (fig. S1c), moving from 10.75 meters to 2.75 meters in front of the observers. In addition to the ‘hit’ trajectory (hit the eye) and ‘near-miss’ trajectory (passing at 5 cm of horizontal shift from the eye), a ‘receding’ trajectory was also included with the reverse trajectory as in the ‘hit’ condition. For the 7T experiment, the on-screen size of the object changed between 0.32 and 1.25 degrees of visual angle, at about 3.99 degrees of eccentricity (angle with vertical meridian is 60º). For the 3T experiment, the on-screen size of the object changed between 0.45 and 1.75 degrees of visual angle, at about 5.58 degrees of eccentricity (angle with vertical meridian is 60º). Compared with the stimulus in experiment 2, the looming stimulus was smaller and disappeared earlier (time-to-collision at the vanishing point was 115 ms). Stimuli were presented either in the normal visual field or the blind visual field of hemianopic patients. In principle, the stimulus location was selected based on the scotoma of each patient. Stimuli were presented at the lower visual field for P02 and P08, and at the upper visual field for other patients. For P11, stimuli were presented in the upper quadrant of the BVF, while in the lower quadrant of the NVF because he could see better than in the upper quadrant. For P12, visual stimuli were presented at an eccentricity of 7.84 degrees, and much closer to the vertical meridian (angle with vertical meridian is 30º). Since the stimulus location of P12 was very different from those of other subjects, we didn’t use his data to generate the group-averaged results in fig. 6.

During fMRI scans, patients were asked to keep fixation and to detect occasional color changes of the fixation point. 6 runs were collected for each patient, except 5 runs for P01 and 4 runs for P03. Each run consisted of 24 trials in which each trajectory was repeated 4 times in the NVF and 4 times in the BVF. The ISI was randomly selected from 8, 10, 12 and 14 seconds.

Four patients (P08∼P12) also performed a behavioral test of stimulus visibility in their BVF. In the subjective visibility test, they were asked to keep fixation and report the visibility to stimuli presented to the NVF or BVF. In the 2-IFC test, the stimulus was presented in one of two 330-ms intervals, separated by a 330-ms gap in-between. The first interval was accompanied by a low-pitch tone at 300 Hz and the second by a high-pitch tone at 700 Hz. Patients were required to keep fixation and to report in which interval the object was presented. Auditory feedback was given after incorrect answers. 240 trials were collected for each patient, including 24 trials in the NVF (8 trials for each trajectory) and 216 trials in the BVF (72 trials for each trajectory). In the behavioral test, the looming stimulus was very close to that in the 7T experiment, with an on-screen size expanding approximately from 0.31 to 1.21 degrees at 3.95 degrees of eccentricity. For P12, the stimulus was at 7.84 degrees of eccentricity.

#### MRI data acquisition and analysis

MRI data for P01, P04, P05, P07, P10 and P11 were acquired with the same 7T scanner, head coil and pulse sequences as in experiment 2. For P01, functional images were acquired with GE-EPI at a higher spatial resolution (1.2-mm isotropic voxels, 62 axial slices of 1.2-mm thickness, 150×150 matrix, TR/TE = 2000/22 ms, nominal flip angle = 78°, partial Fourier factor = 6/8, GRAPPA acceleration factor = 2, multiband factor = 2, bandwidth = 1587 Hz/Px, phase encoding direction from A to P).

P02, P03, P06, P08, P09 and P12 had nonmagnetic metal implants. Due to safety issues of overheating at ultrahigh magnetic fields, their data were acquired with a 3T scanner (Siemens Prisma, Erlangen, Germany) using a 20-channel phased array coil (Nova Medical, Cambridge, MA, USA). High-resolution anatomical images were acquired using a T1-weighted MPRAGE sequence (1-mm isotropic voxels, 192 sagittal slices at 1-mm thickness, image matrix = 256 × 224, TR/TE = 2600/3.02 ms, inversion time = 900 ms, flip angle = 8°, bandwidth = 130 Hz/Px, phase partial Fourier = 6/8, slice partial Fourier = 7/8, no in-plane acceleration). Functional images were acquired with a GE-EPI sequence (2-mm isotropic voxels, 52 or 54 axial slices of 2-mm thickness, 96 × 96 matrix, FOV = 192 × 192 mm, TR = 2000ms, TE =30 or 31.4 ms, flip angle = 80°, multiband or SMS factor = 2, partial Fourier factor = 7/8 or none, bandwidth = 1860 or 2170 Hz/Px, phase encoding direction from A to P, no in-plane acceleration). A few EPI images with reversed phase encoding direction (P to A) were acquired to correct image distortions in the phase encoding direction.

Data analysis procedures were the same as those in experiment 2, except for cortical data analysis. A 6-mm FWHM spatial smoothing was performed on functional volumes after motion correction, followed by per-run scaling and GLM analysis. Statistical volumes were spatially normalized to the MNI space with ICBM 152 symmetric MNI template (2009c) with a combination of linear and nonlinear transformations. Group-level statistics were generated in the MNI space, and then projected to the standard cortical surface (std141) as shown in fig. 6b.

Since the stimulus location of P12 was very different from those of other subjects, we didn’t use his data to generate the group-averaged results for the subcortical area with strong retinotopic representations, including the SC, pulvinar and LGN. But P12 was included in the individual results in fig. 6i and 6j. P03 had severe damage and distortion in his ipsilesional visual thalamus, thus was not included in the results for ipsilesional pulvinar and LGN. Due to a similar reason, P09 was not included in the analysis of ipsilesional LGN. Therefore, for the group-level analysis, there were 11 contra- and ipsi-lesional SCs, 11 contralesional pulvinars and 10 ipsilesional pulvinars, 11 contralesional LGNs and 9 ipsilesional LGNs, and 12 contra- and ipsi-lesional VTAs.

### Statistical analysis of the study

#### Behavioral statistics

A permutation test (exact test, in which all possible permutations were considered) was used to infer the significance of difference between discrimination sensitivities in the upper and lower visual field in fig. 1c. A cluster-based permutation test (10000 times of permutations) was used to calculate the significant time periods in fig. 1d (Nichols & Holmes, 2002). The length of continuous periods (i.e., clusters) of time points showing significant difference in each permutation was used to control the family wise error. Paired t-tests were used to calculate the difference in pupil diameters between different trajectories in fig. 1e and 1f. The difference of behavioral performance of hemianopic patients against chance-level and between trajectories were assessed using a binomial generalized linear mixed model with a logit link function. In the model we tested a fixed effect of trajectory while controlling for a random effect of subjects, with likelihood ratio tests method. The effect of SC responses on the behavioral performance was examined via a linear mixed model analysis, with random effect of subjects.

#### Statistical maps of fMRI

Standard random-effects analysis was used to generate the statistical maps of subcortical and cortical areas. To control the family-wise error of statistical maps within each subcortical ROI (small volume correction), a cluster-mass based permutation test was used to calculate the p value of significant clusters (Hayasaka & Nichols 2004). The cluster mass was defined as the absolute sum of T values in a cluster. The cluster defining threshold is voxel’s p < 0.05. In each permutation, the hit and miss conditions were randomly switched for each subject, followed by a group-level t-test across subjects to generate the statistical map. The largest cluster mass was then recorded for this permutation. The permutation procedure was repeated 10000 times to get the null distribution for the largest cluster mass. A one-sided p value for the actual cluster size was then derived from the null distribution. To obtain accurate null distribution, we subtracted the group-averaged map from each individual map (demean) and removed permutations less than 15% or 85% of data exchanges. Family-wise errors of surface clusters (at p < 0.05 for individual vertex) were determined by Monte Carlo simulation in AFNI.

#### Statistical analysis of ROI-averaged responses

Statistical analysis of ROI-averaged responses was performed with two-way repeated measures ANOVA. To control family-wise errors, we followed the Fisher logic (Levin 1994) and only performed post-hoc t-tests when there was a significant main effect or interaction of the two-way ANOVA. Associations between variables are assessed with Pearson or Spearman correlations. We used Spearman correlation when data pairs contained outliers detected by the Mahalanobis method, and used Pearson correlation otherwise. The false discovery rates of collision sensitivity across ROIs in fig. 4a were calculated by the Benjamini-Hochberg method. The family-wise errors of the correlations in fig. 5a were calculated by a permutation test. The correspondences of data pairs were randomly shuffled in each permutation. The largest absolute value of correlation coefficients was then recorded to compose the null distribution. The corrected p values were derived from the null distribution for the largest correlation coefficient.

## Data and code availability

All data for healthy participants and code from this study have been uploaded to OpenNeuro.org and will be published with the study. Patient data are not publicly available but can be requested from Peng Zhang.

Note: Supplementary information is available for this preprint.

