## Supplementary information for "Human subcortical pathways automatically detect collision trajectory without attention and awareness"

### Supplemental materials

#### a: Experiment 1

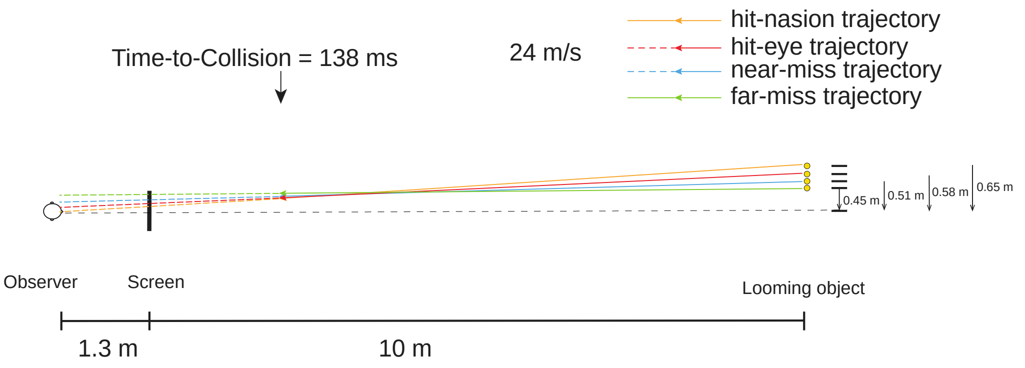


#### b: Experiment 2

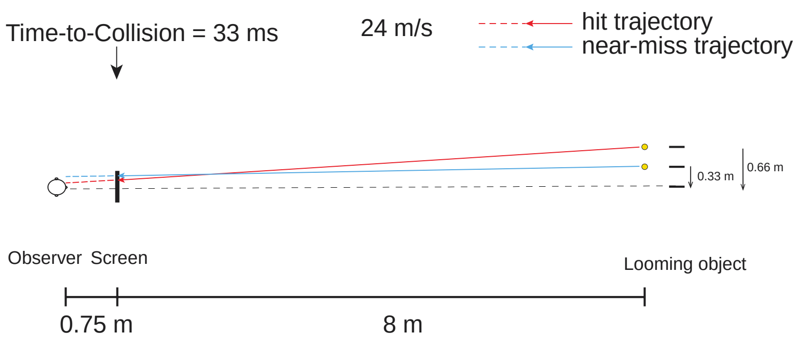


#### c: Experiment 3

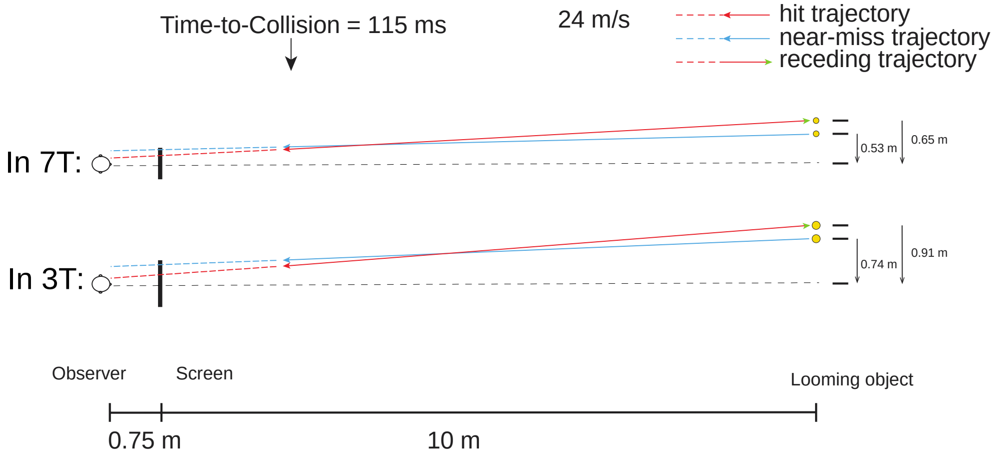

**Figure S1. Top view of stimulus trajectories. (a)** Experiment 1. A baseball sized sphere launched 11.3 meters away from the observer, moving at a speed of 24 m/s. It disappeared at 3.3 meters from the observers at 138 ms of time-to-collision, as indicated by the location of the arrows. Yellow, red, blue and green arrows indicate the hit-nasion, hit-eye, near-miss and far-miss trajectories. Stimuli were presented on a 3D monitor 1.3 meters in front of the observers. (**b)** Experiment 2. Stimuli were presented on a translucent screen with a 2D projector. The sphere moved from 8.75 m away and disappeared on the screen 0.75 m in front of the observer at 33 ms of time-to-collision. Red and blue arrows indicate the hit and near-miss trajectories. **(c)** Experiment 3. In hit and near-miss conditions, the sphere moved from 10.75 m to 2.75 m in front of the observer. The time-to-collision at disappearance was 115 ms. The receding trajectory was the reverse of the hit trajectory. Red, blue and green arrows indicate the hit, near-miss and receding trajectories. The eccentricity and the size of the sphere (3T: 8.4cm; 7T: 6cm) was slightly different in the 3T and 7T scanning.

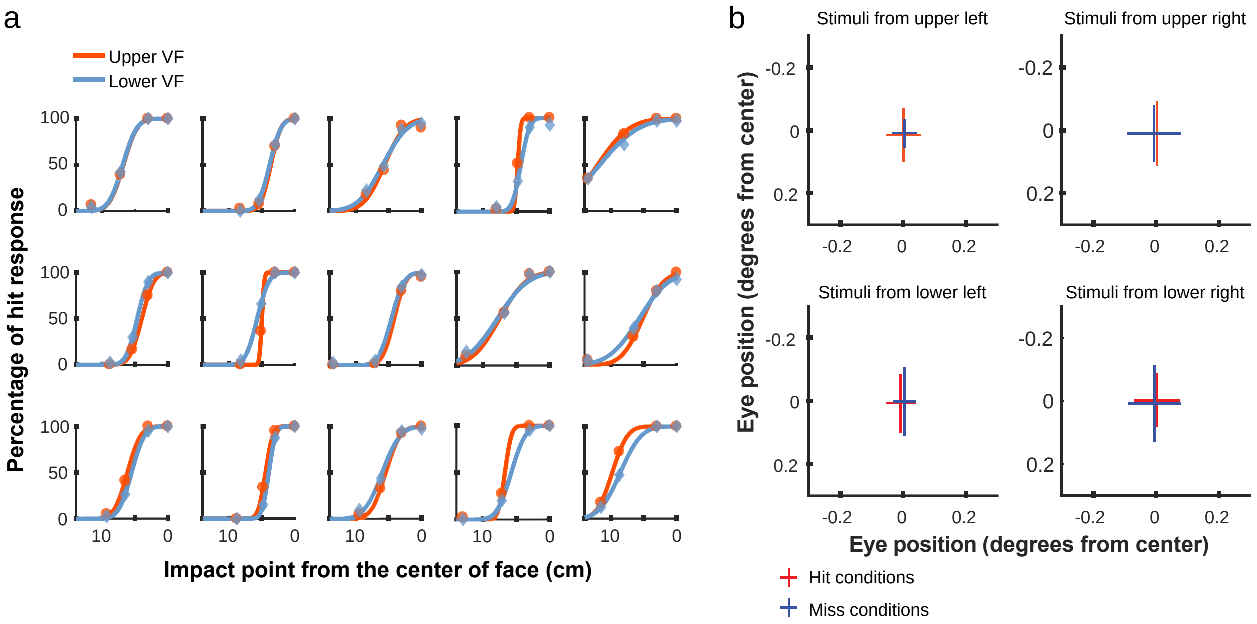


**Figure S2. (a) Individual data for the behavioral experiment (exp. 1).** The percentage of hit responses to looming stimuli as a function of impact points was fitted with a normal CDF. **(b) Distribution of eye positions for hit and miss looming stimuli from four quadrants of the visual field.** The eye gaze positions from stimulus onset to 1000 ms after were analyzed. Error bars indicate 3´standard deviation of the distribution. No significant difference was found between the eye positions to the hit and miss stimuli.

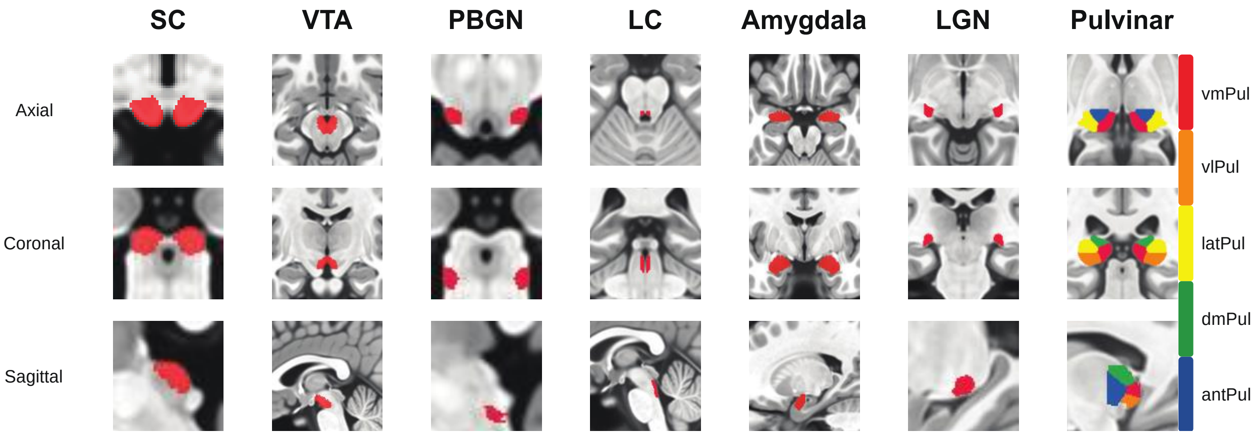


**Figure S3. Anatomical ROIs of subcortical nuclei (exp. 2 and exp. 3).** From left to right are the superior colliculus (SC), ventral tegmental area (VTA), parabigeminal nucleus (PBGN), locus coeruleus (LC), amygdala, lateral geniculate nucleus (LGN), and Pulvinar. The pulvinar was parcellated into 5 subdivisions based on task-coactivation patterns (Barron 2015), including the ventromedial pulvinar (vmPul, red), ventrolateral pulvinar (vlPul, orange), lateral pulvinar (latPul, yellow), dorsomedial pulvinar (dmPul, green) and anterior pulvinar (antPul, blue).

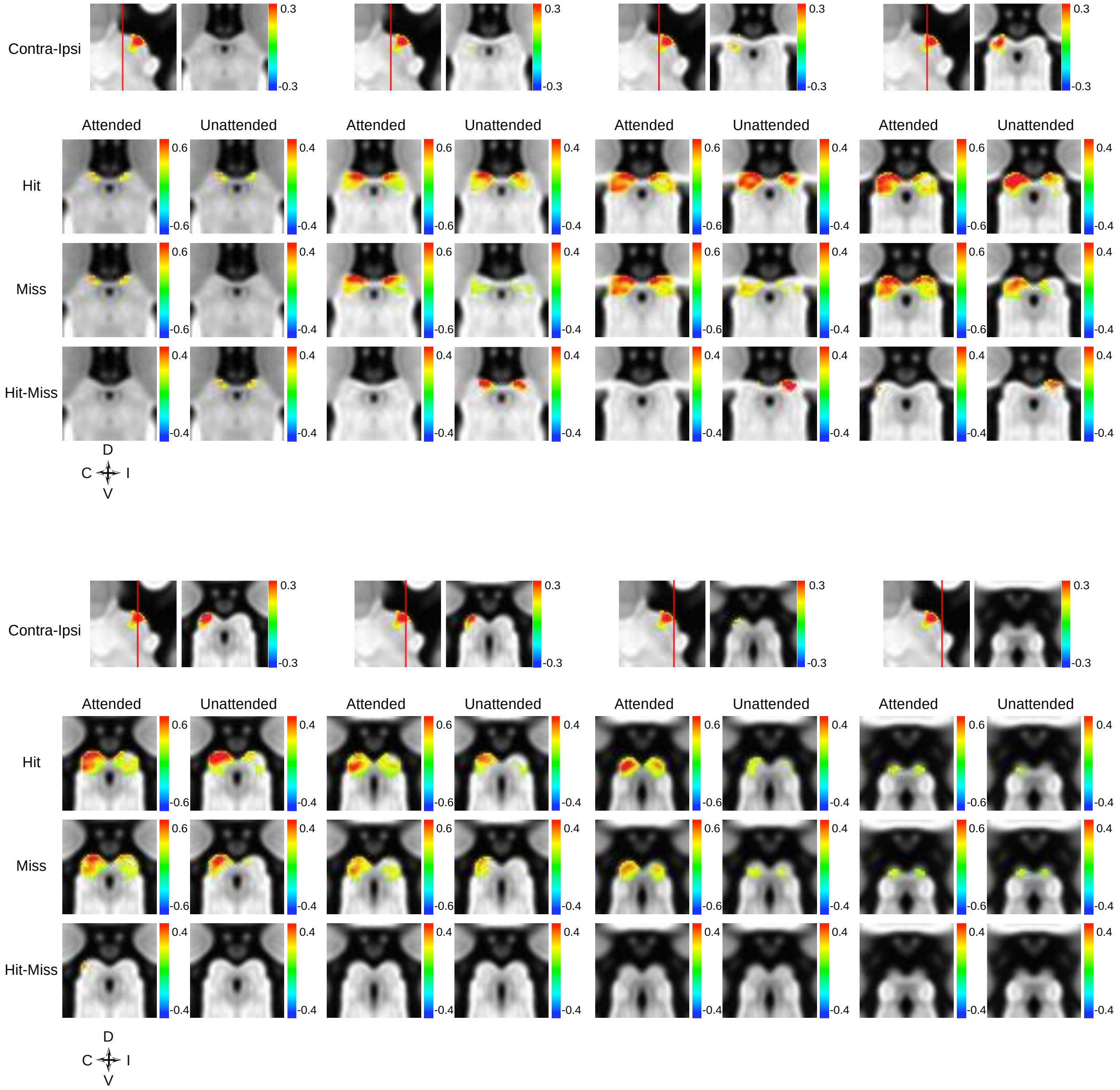


**Figure S4. Looming-evoked responses across the SC (exp. 2).** The first row of each panel shows the retinotopic activations with significantly stronger responses to contralateral than to ipsilateral stimuli. Red lines on the sagittal view indicate the location of the coronal slices. The second to the fourth rows show the activation maps for the Hit, Miss and Hit-Miss responses. Maps were thresholded at voxel $p < 0.05$ uncorrected.

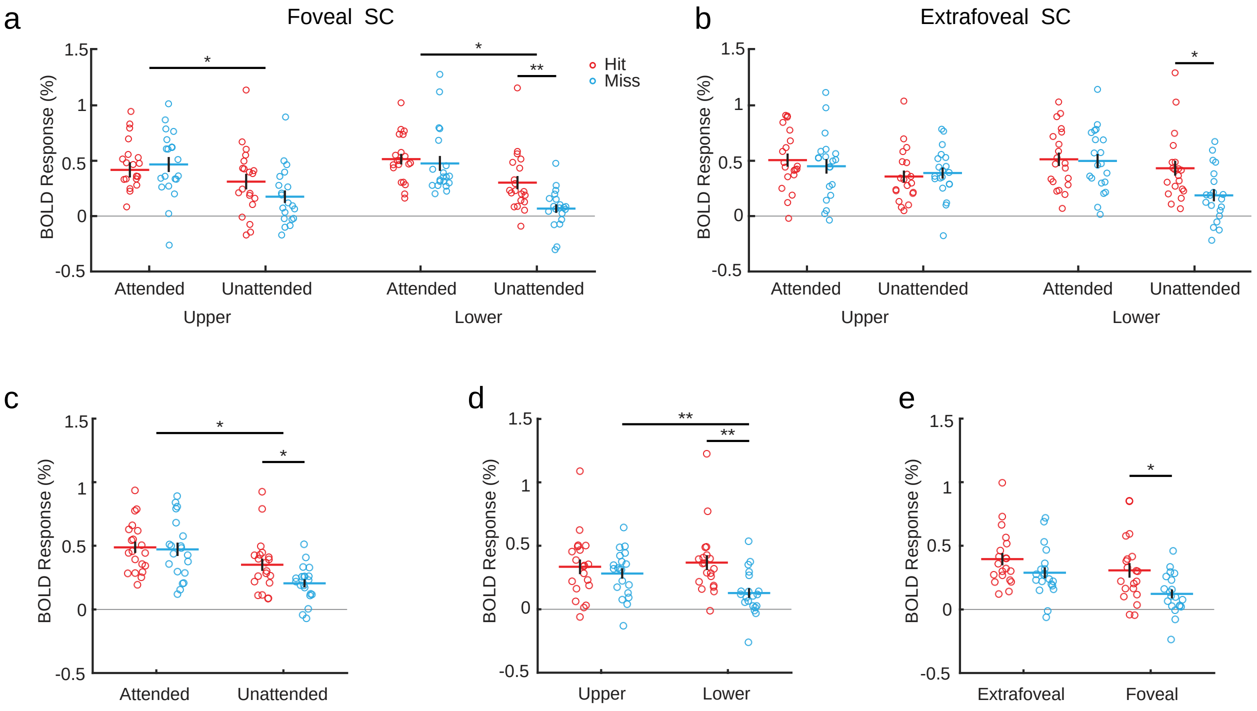


**Figure S5. (a) Responses in the foveal SC to looming stimuli from the upper and lower fields (exp. 2).** For both upper and lower visual fields, there were significant two-way interactions of attention and trajectory (upper: F(1,19) = 4.77, p = 0.042, $\eta_p^2$ = 0.201; lower: F(1,19) = 6.25, p = 0.022, $\eta_p^2$ = 0.247). For the lower visual field, the hit stimulus also evoked significantly stronger response than that of the miss stimulus (t(19) = 3.817, p = 0.002 Bonferroni corrected, Cohen's d = 0.854). **(b) Upper and lower visual field responses in the extrafoveal SC.** For looming stimulus approaching from the lower visual field in the unattended condition, there was a significantly stronger response to hit compared to miss trajectories (t(19) = 2.770, p = 0.024 corrected, Cohen's d = 0.619). **(c) Looming-evoked responses in the attended and unattended conditions.** Responses were averaged across foveal/extrafoveal SCs and upper/lower visual fields. There were significant effects of attention (F(1,19) = 30.26, p < 0.001, $\eta_p^2$ = 0.614), trajectories (F(1,19) = 6.94, p = 0.016, $\eta_p^2$ = 0.268) and interaction (F(1,19) = 5.29, p = 0.033, $\eta_p^2$ = 0.218). Post-hoc paired t-tests revealed a significant collision sensitivity in the unattended condition (t(19) = 2.986, p = 0.016 corrected, Cohen's d = 0.668), but not in the attended condition (t(19) = 0.464, p = 0.648 uncorrected, Cohen's d = 0.104). These results demonstrate strong attention modulations on looming-evoked responses, which almost saturated the SC's response in the attended condition. **(d) Looming-evoked responses to stimulus from the upper and lower visual fields.** Since responses were almost saturated in the attended condition, only results from the unattended condition were shown here, averaged across foveal and extrafoveal SCs. There was a significant collision sensitivity to stimulus in the lower visual field (t(19) = 3.626, p = 0.004 corrected, Cohen's d = 0.811), but not in the upper visual field (p = 0.455 uncorrected). Also looming stimuli on a near-miss trajectory from the upper visual field evoked significantly stronger response in the SC compared with those from the lower visual field (t(19) = 3.204, p < 0.01 corrected, Cohen's d = 0.716). **(e) Looming-evoked responses in the foveal**

**and extrafoveal SCs.** Only results from the unattended condition were shown. Extrafoveal SC evoked significantly stronger responses than the foveal SC ($F(1,19) = 18.42$, $p < 0.001$, $\eta_p^2 = 0.492$), a significant collision-sensitive response can be observed in the foveal SC ($t(19) = 3.129$, $p = 0.012$ corrected, Cohen's $d = 0.700$), but not in the extrafoveal SC ($p = 0.094$ uncorrected).

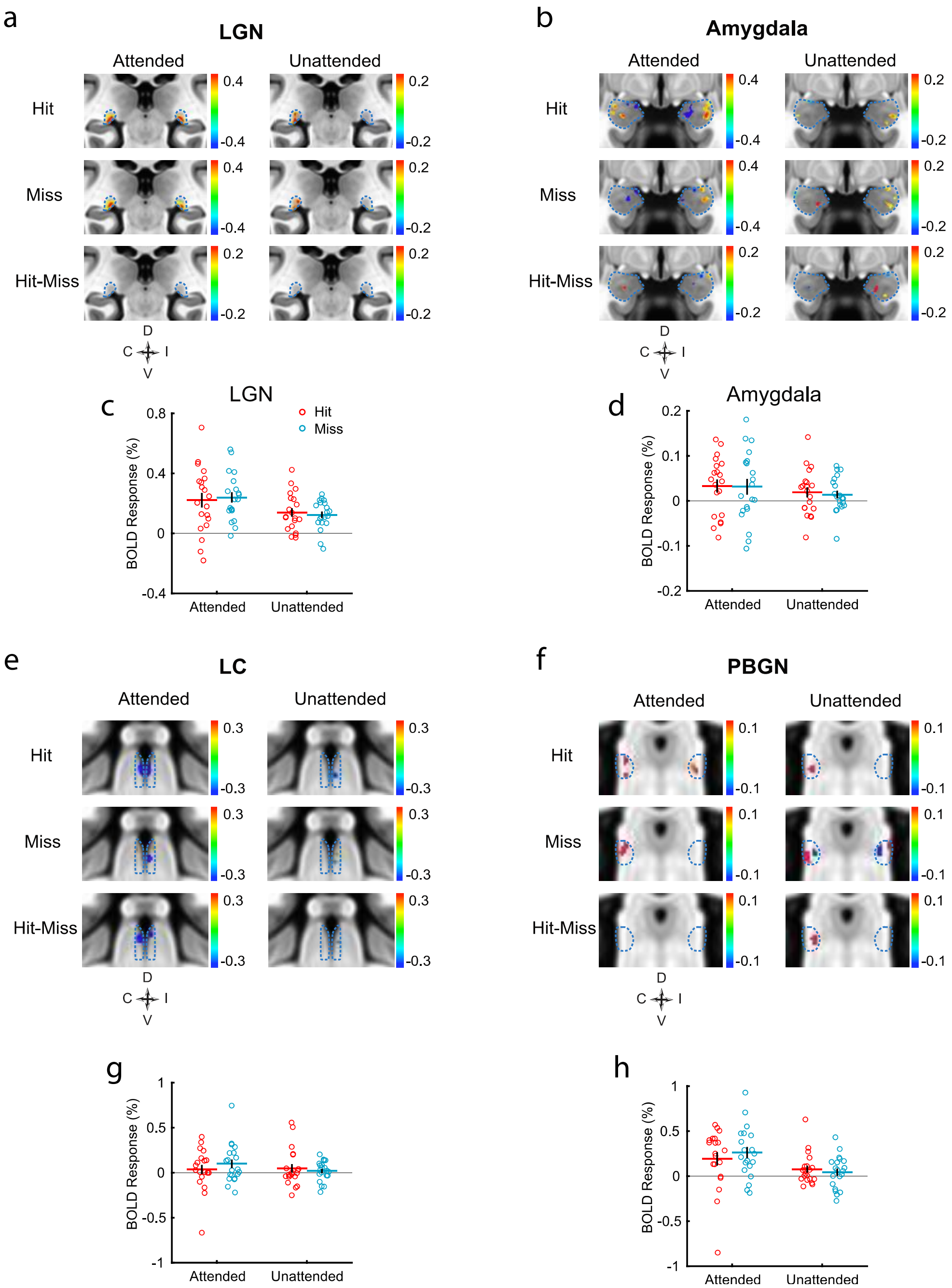


**Figure S6. Activation maps and ROI-averaged responses to looming stimuli in other subcortical nuclei (exp. 2). (a,c)** LGN. **(b,d)** Amygdala. **(e,g)** LC. **(f,h)** PBGN. No significant collision-sensitive cluster or ROI-averaged response can be found from these areas. Maps were thresholded at $p < 0.05$ uncorrected.

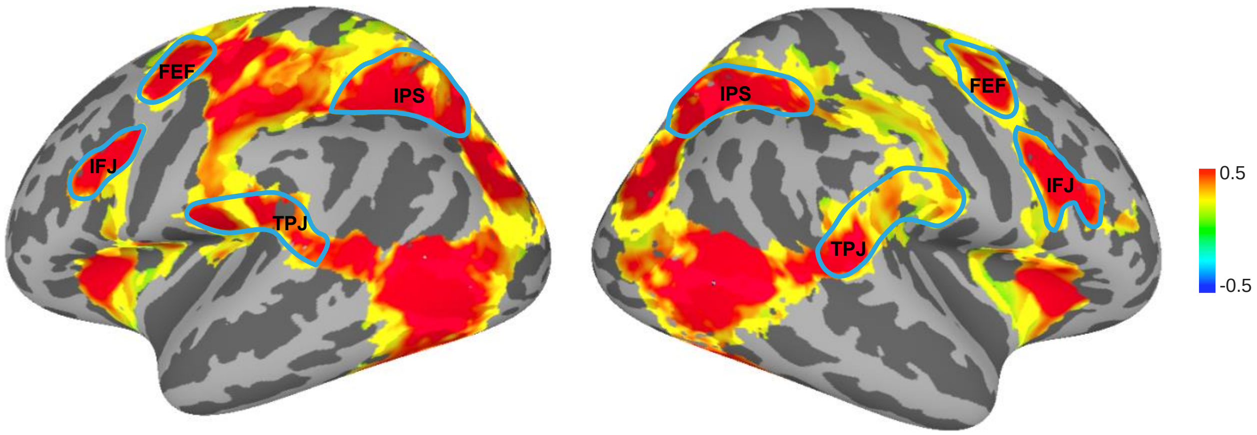


**Figure S7. ROI definitions for frontoparietal attention networks in the correlation analysis (exp. 2).** The activation map shows the group-averaged activation across all stimulus conditions (thresholded at $p < 0.001$ uncorrected). Blue lines indicate the boundaries of the ROIs.

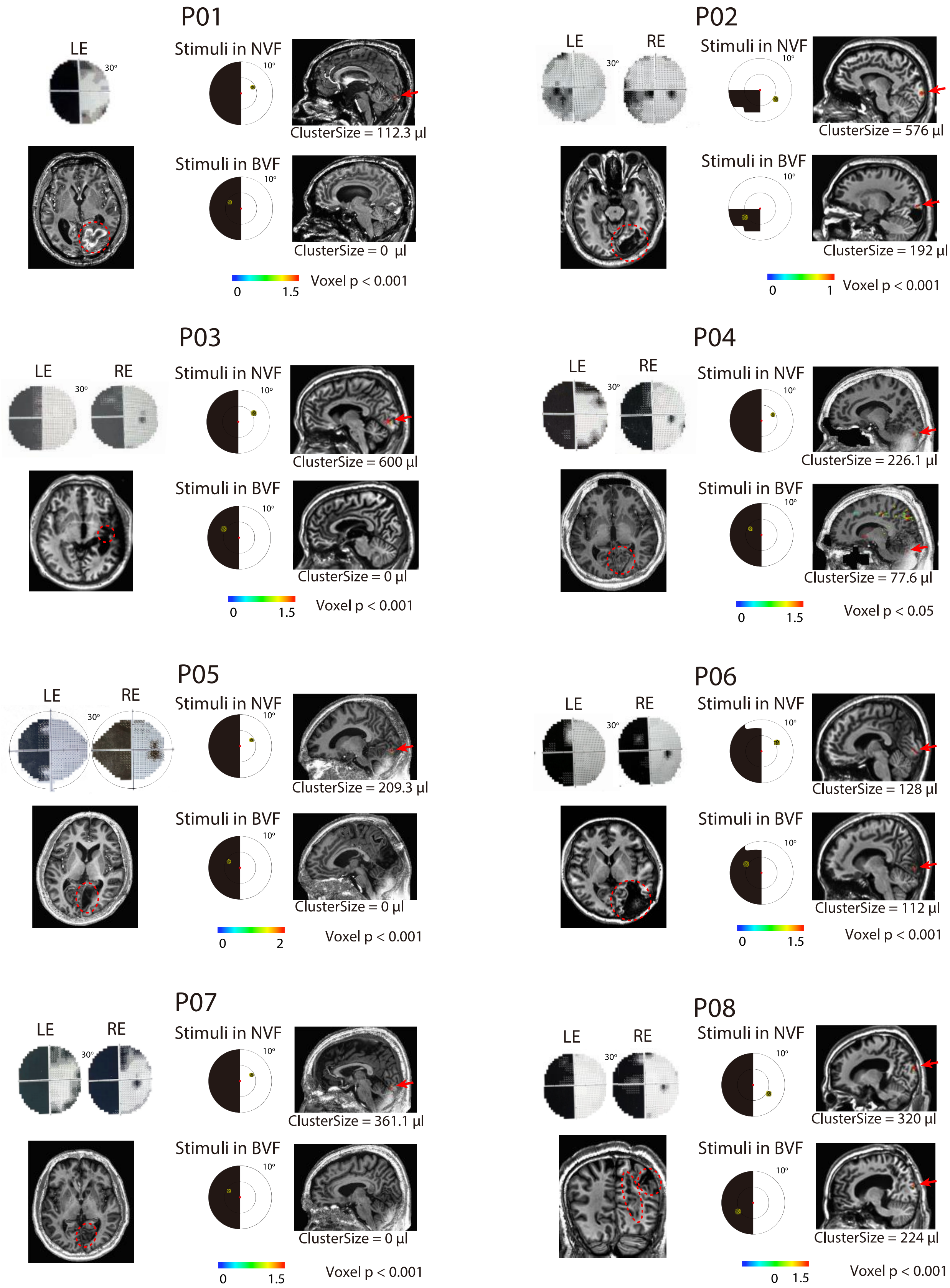
P01
LE
30°
Stimuli in NVF
10°
ClusterSize = 112.3 µl
Stimuli in BVF
10°
ClusterSize = 0 µl
0
1.5
Voxel p < 0.001
P02
LE
RE
30°
Stimuli in NVF
10°
ClusterSize = 576 µl
Stimuli in BVF
10°
ClusterSize = 192 µl
0
1
Voxel p < 0.001
P03
LE
RE
30°
Stimuli in NVF
10°
ClusterSize = 600 µl
Stimuli in BVF
10°
ClusterSize = 0 µl
0
1.5
Voxel p < 0.001
P04
LE
RE
30°
Stimuli in NVF
10°
ClusterSize = 226.1 µl
Stimuli in BVF
10°
ClusterSize = 77.6 µl
0
1.5
Voxel p < 0.05
P05
LE
RE
30°
Stimuli in NVF
10°
ClusterSize = 209.3 µl
Stimuli in BVF
10°
ClusterSize = 0 µl
0
2
Voxel p < 0.001
P06
LE
RE
30°
Stimuli in NVF
10°
ClusterSize = 128 µl
Stimuli in BVF
10°
ClusterSize = 112 µl
0
1.5
Voxel p < 0.001
P07
LE
RE
30°
Stimuli in NVF
10°
ClusterSize = 361.1 µl
Stimuli in BVF
10°
ClusterSize = 0 µl
0
1.5
Voxel p < 0.001
P08
LE
RE
30°
Stimuli in NVF
10°
ClusterSize = 320 µl
Stimuli in BVF
10°
ClusterSize = 224 µl
0
1.5
Voxel p < 0.001

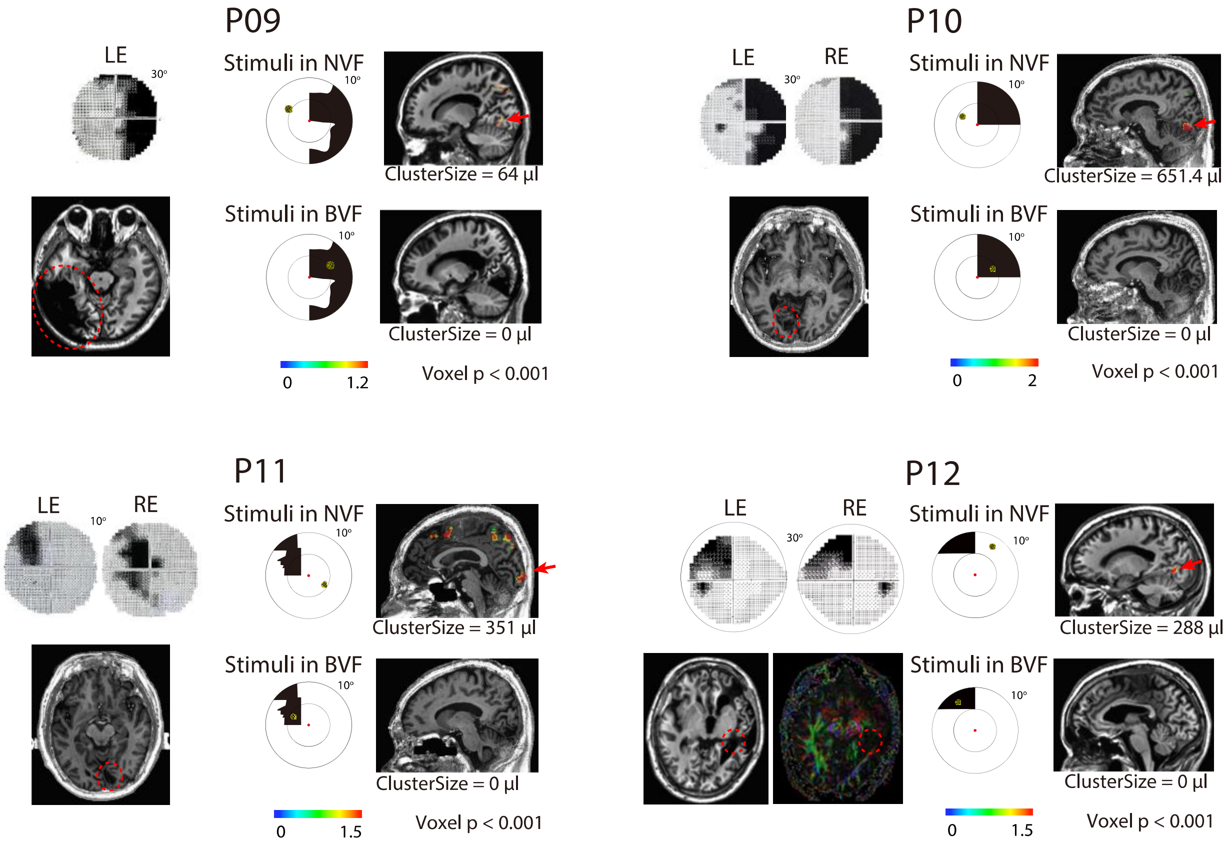


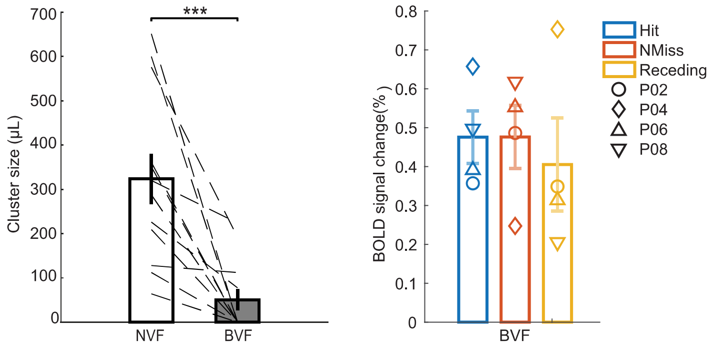


**Figure S8. Clinical perimetry, lesioned locations and stimulus-evoked occipital activations for hemianopic patients (exp. 3).** For each patient, the left panels show the Humphrey perimetry of visual field test. In the structural image below, relevant lesioned locations were indicated by red dashed ovals. For P17, both T1w and diffusion tensor images were shown to indicate the lesion of right optic radiation. In the middle panels, the scotoma was depicted schematically within 10 degrees of eccentricity in black color (i.e., relative sensitivity < -20dB and p < 0.5% compared with normal population), with the yellow sphere indicating the stimulus in the fMRI experiment. The right panels show the occipital activations to stimuli presented to the NVF and BVF (indicated by red arrows).

Although clear contralateral V1 activations can be observed to stimuli presented to the NVF, most patients (8/12) showed no significant V1 activation in the lesioned hemisphere to stimulus presented to the BVF (the left bar graph below: dashed line for individual data, *** for $p < 0.001$). For the four patients (P02, P04, P06 and P08) showing weak uncorrected activations in the occipital lobe of the lesioned hemisphere, no significant difference was found in the responses of these voxels to the hit and miss stimuli (the right bar graph below), which cannot explain the collision-sensitive responses in the SC (fig. 6a).

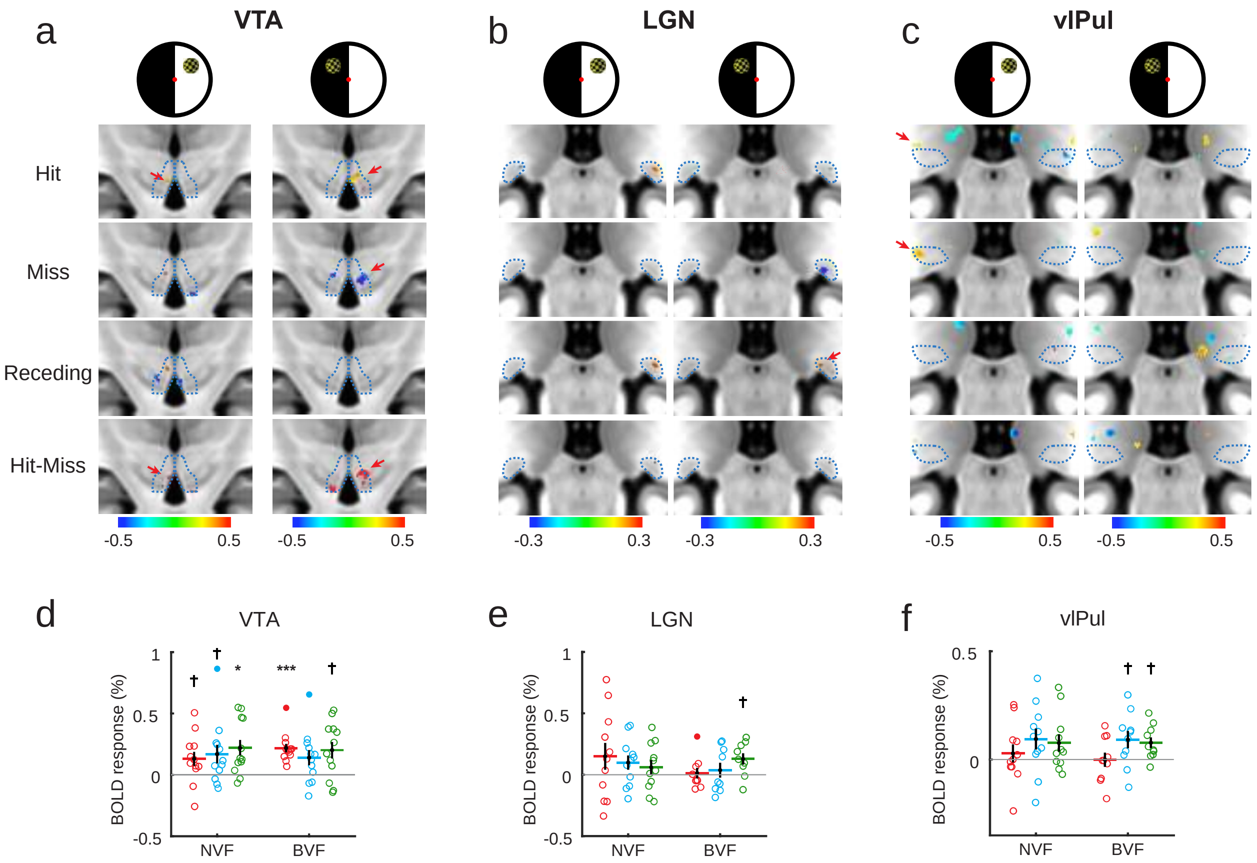


**Figure S9. Looming-evoked responses in the VTA, LGN and vlPul of hemianopic patients (exp. 3). (a-c)** The group-averaged activation maps (thresholded at $p < 0.05$ uncorrected). A significant cluster with collision sensitivity was found in the ipsilesional VTA when stimuli were presented to the BVF (cluster's volume = 37 µl, cluster's $p = 0.047$). Red arrows indicate the location of collision sensitive clusters in the VTA (a), visually evoked response to receding stimuli in the LGN (b), and visually evoked response to hit and near-miss stimuli in the vlPul (c). Blue dotted lines indicate the boundary of ROIs. **(d-e)** ROI-averaged responses of most responsive voxels. * and *** denote $p < 0.05$ and $p < 0.001$ after Holm correction, and † for uncorrected $p < 0.05$. Error bars represent standard errors of the mean.

**Table S1. Clinical characteristics of hemianopic patients**

| Patient | Sex | Age （year） | Lesioned location | Reason | Time post-lesion （month） | Blind visual field |
|---|---|---|---|---|---|---|
| P01 | male | 38 | Right occipital | Tumor surgery | 3 | Left |
| P02 | male | 53 | Right occipital | Cerebral hemorrhage | 11 | Lower Left |
| P03 | male | 36 | Right optic radiation | Cerebral hemorrhage | 1 | Left |
| P04 | male | 37 | Right occipital | Cerebral infarction | 1 | Left |
| P05 | male | 25 | Right occipital, parietal | Hematoma, cerebral hemorrhage | 2 | Left |
| P06 | femal e | 23 | Right occipital, temporal | Cerebral hemorrhage | 5 | Left |
| P07 | male | 63 | Right occipital | Cerebral infarction | 1 | Left |
| P08 | male | 42 | Right occipital, parietal | Brain trauma, hematoma | 9 | Left |
| P09 | male | 60 | Left occipital, temporal | Cerebral hemorrhage | 104 | Right |
| P10 | male | 57 | Left occipital | Cerebral infarction | 2 | Right |
| P11 | male | 64 | Right occipital | Cerebral infarction | 16 | Upper left |
| P12 | male | 48 | Right optic radiation | Brain trauma, cerebral hemorrhage | 280 | Upper left |
